# Live-cell imaging reveals the spatiotemporal organization of endogenous RNA polymerase II phosphorylation at a single gene

**DOI:** 10.1101/2020.04.03.024414

**Authors:** Linda S. Forero-Quintero, William Raymond, Tetsuya Handa, Matthew Saxton, Tatsuya Morisaki, Hiroshi Kimura, Edouard Bertrand, Brian Munsky, Timothy J. Stasevich

## Abstract

The carboxyl-terminal domain of RNA polymerase II is dynamically phosphorylated during transcription in eukaryotic cells. While residue-specific phosphorylation has been mapped with exquisite spatial resolution along the 1D genome in a population of fixed cells using immunoprecipitation-based assays, the timing, kinetics, and spatial organization of phosphorylation along a single-copy gene have not yet been measured in living cells. Here, we achieve this by combining multi-color, single-molecule microscopy with fluorescent antibody-based probes that specifically bind to unphosphorylated and phosphorylated forms of endogenous RNAP2 in living cells. Applying this methodology to a single-copy HIV-1 reporter gene provides live-cell evidence for heterogeneity in the distribution of RNAP2 along the length of the gene as well as clusters of Serine 5 phosphorylated RNAP2 that form around active genes and are separated in both space and time from nascent mRNA synthesis. Computational models fit to our data determine that 5 to 40 RNAP2 cluster around the promoter of a gene during typical transcriptional bursts. Rapid imaging demonstrates nearly all RNAP2 in the cluster acquire Serine 5 phosphorylation within 3-6 seconds of arrival. Transcription from the cluster appears to be highly efficient, with nearly half of the clustered RNAP2 ultimately escaping the promoter in 1.5 minutes on average to elongate a full-length mRNA in approximately five minutes. The highly dynamic and spatially organized concentrations of RNAP2 we observe support the notion of highly efficient transcription clusters that form around promoters and contain high concentrations of RNAP2 phosphorylated at Serine 5.

---

In eukaryotic cells, the catalytic RPB1 subunit of RNA polymerase II (RNAP2) possesses an extended carboxy terminal domain (CTD) that consists of heptapeptide repeats (52 in humans) with a consensus sequence (Tyr_1_-Ser_2_-Pro_3_-Thr_4_-Ser_5_-Pro_6_-Ser_7_). The CTD region is dynamically phosphorylated as RNAP2 progresses through the transcription cycle, regulating each step of transcription, from initiation to termination. In some models, RNAP2 is recruited to promoters in an unphosphorylated form (CTD-RNAP2), but is later phosphorylated at Serine 5 (Ser5ph-RNAP2) upon initiation and at Serine 2 (Ser2ph-RNAP2) during active elongation^1–4^. Interest in the CTD has recently increased due to observations of highly dynamic RNAP2 clustering^4‒6^ that correlates with the phosphorylation status of the CTD^7,8^. In particular, recent data suggest that a transcriptional cluster forms around gene promoters early in the transcription cycle. The cluster is thought to be enriched in unphosphorylated- and Ser5ph-RNAP2 that appear to constrain chromatin movement near the transcription start site^9^. However, upon transcriptional activation, hyperphosphorylation of RNAP2 at Ser2 allows the enzyme to escape the cluster and begin active elongation^7,9^. The dynamic clustering of RNAP2 involves many steps and a complex orchestration of multiple factors and could therefore represent a global form of transcriptional regulation^10^.

RNAP2 phosphorylation throughout the transcription cycle has traditionally been studied in fixed cells using immunoprecipitation-based assays^1,3,11^. These studies provide precise spatial maps of the average positions of RNAP2 along the 1D genome. Unfortunately, the inherent averaging masks heterogeneity and the procedure limits temporal resolution to timescales of tens of minutes or longer^12^. RNAP2 dynamics can instead be imaged and quantified in living cells using fluorescence microscopy, overcoming the limitations of traditional assays. Recent singlemolecule tracking technologies^13–17^ have made it possible to monitor single RNAP2 as they bind at non-specific locations throughout the genome^5,18^ as well as at specific, single-copy genes^6,17^ premarked with MS2^19,20^ or PP7^21^ RNA stem loops (that are lit up co-transcriptionally when, respectively, fluorescent MS2 or PP7 coat proteins bind to them). Each of these studies used permanent fluorescent fusion tags to track RNAP2. Fusion tags are incapable of discerning post-translational modifications to RNAP2, including transcription cycle associated phosphorylation events.

One way to resolve post-translational modifications to RNAP2 is to use antibody-based probes that bind and light-up specific modifications to residues within the CTD *in vivo^22–26^.* However, the signal-to-noise is limited with this approach because of the presence of unbound and freely diffusing probes that increase the fluorescence background. Applications have therefore been restricted to large tandem gene arrays. Signal-to-noise is amplified by the multiple copies of a gene within these arrays, but heterogeneity from one gene copy to another is again masked by averaging^27^. Therefore, the spatiotemporal dynamics of RNAP2 phosphorylation at single-copy genes remain unclear.

Here we combine multi-color single-molecule microscopy, complementary fluorescent antibodybased probes, and rigorous computational modeling to visualize, quantify, and predict endogenous RNAP2 phosphorylation dynamics at a single-copy reporter gene in living cells. This unique combination of technologies allows us to directly visualize the temporal ordering and spatial organization of RNAP2 phosphorylation and mRNA synthesis throughout the transcription cycle at the reporter gene. We find evidence for relatively high concentrations of RNAP2 near the beginning versus end of the gene that are both spatially and temporally separate from elongating RNAP2 and nascent mRNA synthesis. Collectively, our data provide live-cell support for the existence of higher-order, phosphorylation-dependent transcriptional clusters that dynamically form and surround active genes throughout the transcription cycle.

## Results

### Technology to visualize endogenous RNAP2 transcription cycle dynamics at a single gene

To visualize the spatiotemporal dynamics of endogenous RNAP2 phosphorylation at a single gene, we used an established HeLa cell line (H-128) harboring an MS2-tagged HIV-1 reporter gene and stably expressing both GFP-tagged MS2 coat protein (MCP) and an untagged HIV-1 trans-activator of transcription (Tat)^20^. We chose HIV-1 as our reporter gene because it is a prototypical model for RNAP2 phosphorylation^28^. The HIV-1 reporter is strongly active in our cell line due to persistent stimulation by Tat, producing a bright MCP signal that pinpoints the location of the transcription site and gauges its activity in real-time^20^ (Fig. 1a). Consistent with this strong signal, immunostaining experiments in fixed cells revealed the transcription site is highly enriched in RNAP2 and relatively depleted in histones and their epigenetic modifications (Sup. Fig. 1a-b). Chromatin immunoprecipitation (ChIP) experiments furthermore confirmed the presence of CTD-RNAP2 and its phosphorylated forms Ser5ph- and Ser2ph-RNAP2, respectively. In particular, we detected that CTD-RNAP2 and Ser5ph-RNAP2 signals are highest at the transcription start site, whereas Ser2ph-RNAP2 is highest towards the end of the gene (Fig. 1b). However, because these data come from a population of fixed cells, whether the various forms of RNAP2 are present at the same time and place and whether or not they appear in a preferred order is difficult to extract from this assay.

**Fig. 1:**
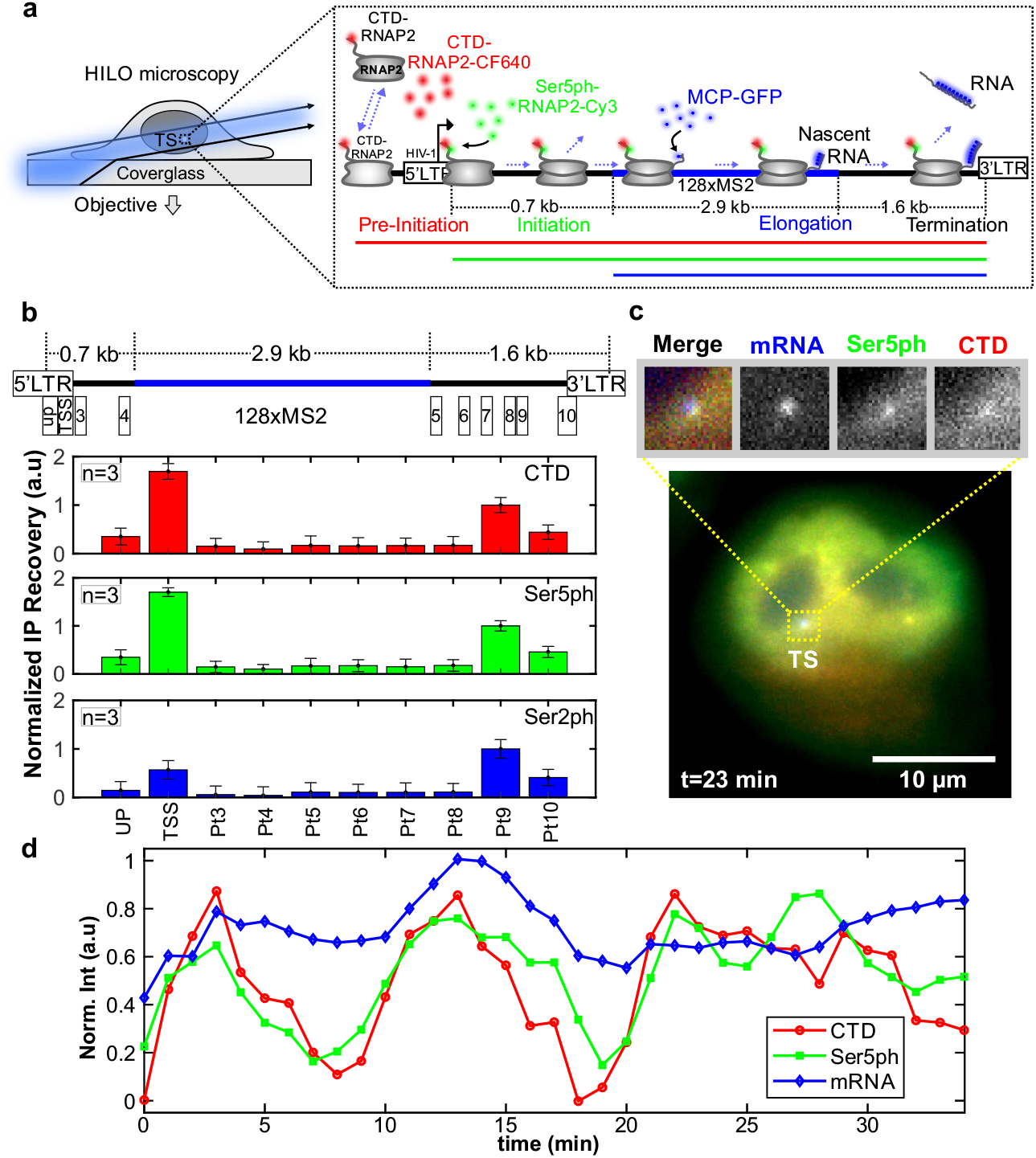
A system for imaging the endogenous RNAP2 transcription cycle at single genes. **(a)** Schematic of the system. The reporter gene is controlled by the HIV-1 promoter and is tagged with a 128xMS2 cassette (blue bar). RNAP2 is represented in gray. RNA is marked by MCP-GFP that binds to the transcribed MS2 stem loops (mRNA, blue). The recruited and initiated RNAP2 are labeled by Fabs (conjugated with CF640 and Cy3) that bind unphosphorylated CTD RNAP2 heptad repeats (CTD, red) and Serine 5 phosphorylated repeats (Ser5ph, green), respectively. **(b)** Average chromatin immunoprecipitation occupancy of CTD-RNAP2 (red, upper panel), Ser5ph-RNAP2 (green, middle panel), and Ser2ph-RNAP2 (blue, lower panel) across the HIV-1 reporter gene (positions 1-10 are highlighted in the cartoon above). **(c)** Sample live cell showing CTD-RNAP2, Ser5ph-RNAP2, and mRNA co-localizing at the transcription site (TS). **(d)** Normalized intensity at the TS over time from the cell in (c) for CTD-RNAP2 (red circles), Ser5ph-RNAP2 (green squares), and mRNA (blue diamonds).

To better characterize the spatiotemporal dynamics of single-cell RNAP2 modifications during transcription, we loaded fluorescent fragmented antibodies (Fab, generated from the same antibodies used in ChIP)^22,29^ recognizing (1) the CTD of RNAP2 (anti-CTD RNAP2) without or with residue-specific phosphorylations, and (2) heptad repeats within the CTD that are phosphorylated at Serine 5 (anti-Ser5ph RNAP2). These antibodies have previously been shown to be specific for their respective targets via Western blotting and ELISA^27^, and in ChIP-seq^30^ experiments. Fab generated from these antibodies have also been shown to rapidly bind and unbind their targets, making them valuable for monitoring temporal changes in RNAP2 phosphorylation^27^. Consistent with anti-CTD Fab labeling all RNAP2 and anti-Ser5ph Fab labeling a subset of RNAP2, we observed regions within the nucleus with RNAP2 enriched or depleted with Ser5ph (Sup. Fig. 2). These capabilities allowed us to distinguish three distinct steps of the transcription cycle at the HIV-1 reporter gene: RNAP2 recruitment (marked by Fab against CTD-RNAP2), initiation (marked by both Fab against CTD-RNAP2 and Fab against Ser5ph-RNAP2), and elongation (marked by all Fab and MCP binding to mRNA), as depicted in Fig. 1a,c. Although we attempted to also visualize Ser2ph at the locus with our Fab, signal-to-noise was insufficient to detect in living cells, presumably because the antibody is not sensitive enough to recognize this modification at the single-gene level.

Nevertheless, this setup has several advantages that collectively enhance signal-to-noise at the transcription site. First, Fab bind endogenous RNAP2, so all RNAP2 in the cell have high likelihood to be labeled without having to genetically engineer a fusion knock-in tag^18,31^ and/or alpha-amanatin resistance^5^. Second, fluorescence is naturally amplified since mammalian RNAP2 contains 52 heptad repeats in its CTD^32^, each of which can be bound by a fluorescent Fab at the transcription site. Third, Fab continually bind and unbind RNAP2, mitigating the loss of fluorescence due to local photobleaching. In combination with a multi-color, single-molecule microscope^33^ employing oblique HILO illumination to enhance signal-to-noise by an order of magnitude^13^, these advantages allowed us to generate movies in which we monitored endogenous RNAP2 phosphorylation dynamics at the HIV-1 reporter gene in 3-colors.

As shown in Fig. 1c-d, movies revealed correlated fluctuations between the mRNA signal and endogenous CTD-RNAP2 and Ser5ph-RNAP2 signals at the transcription site. To ensure correlations were not an artifact of focusing issues, we tracked the transcription site in 3D (by imaging 13 z planes per time point) to keep the MS2 signal continually in focus (Sup. Fig. 1c, left panel). The correlations were also not caused by photobleaching, as signals fluctuated both up and down throughout the entire imaging time course, remaining on average constant (Sup. Fig. 1c). Finally, to rule out the possibility that correlated fluctuations were caused by bleed through from one fluorescence channel to another, we re-imaged cells lacking Fab. In all cases, no bleed through was observed (Sup. Fig. 1d-e), as quantified by the covariance between channels (Sup. Fig. 1f). We therefore conclude the correlations reflect natural bursts in endogenous transcriptional activity at the HIV-1 reporter gene, demonstrating our ability to detect and quantify endogenous RNAP2 phosphorylation dynamics at a single-copy gene.

### Long-term imaging of fluctuations at the reporter gene reveals temporal ordering of RNAP2 phosphorylation

In the majority of cells, the mRNA was steadily produced by the HIV-1 reporter gene, with strong signals persisting for hours at a time. In a few cells the mRNA signal completely disappeared, indicating a loss of nearly all transcription activity. We were interested in capturing these rare events in a single time course to better discriminate the relative timing of our RNAP2 and mRNA signals. To accomplish this, we adjusted our imaging conditions to optimize detection of all three signals in single cells over a period of three hours (200 time points), as exemplified in Fig. 2a-c (also see Sup. Movie 1, and Sup. Fig. 3a-c). We again imaged in z-stacks (13 planes spaced by 0.5 *μ*m) covering the whole nucleus at each time point throughout the entire experiment. We were therefore confident that the fluctuations were due to changes in transcription activity and not related to transcription site movement into and out of the focal plane (Sup. Fig. 3d). With these imaging conditions we found cells in which the mRNA signal turned on and off up to four times, indicating bursts of transcription and multiple complete transcription cycles. Consistent with our previous result, signals at the transcription site were highly correlated and fluctuated generally in unison, although there were distinct periods of time when one signal could be seen for multiple frames in the absence of some other signals. This again ruled out bleed through and suggested the signals were not perfectly synchronized. To ensure the correlated fluctuations were specific to the locus and not cell-wide, we verified that covariances between the mRNA signal and the CTD or Ser5ph signals were significantly stronger when both signals were measured at the transcription site compared to when one or both signals were measured a short distance (p1) from the transcription site (Sup. Fig. 3e-g).

**Fig. 2:**
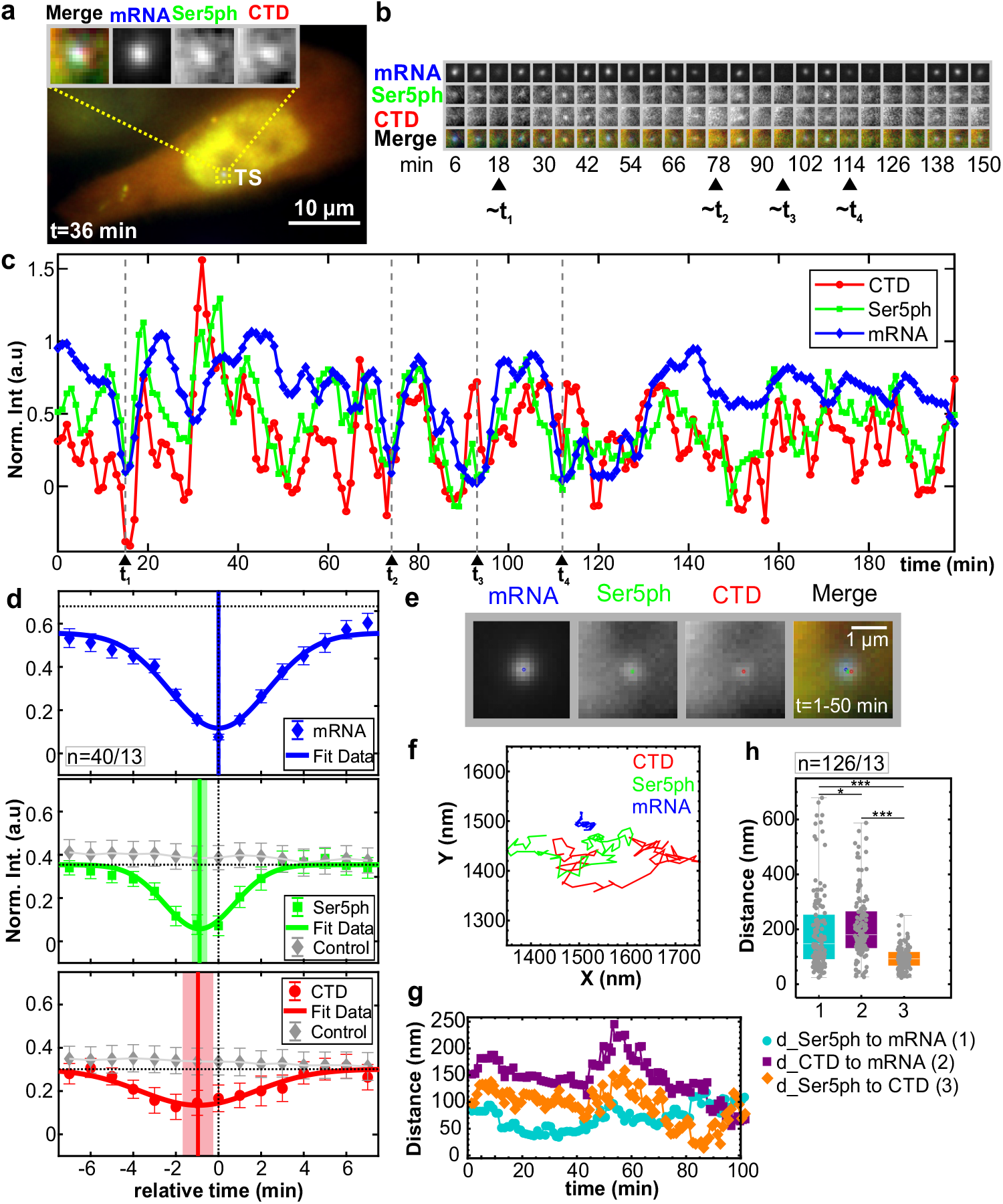
Spatiotemporal organization of the RNAP2 CTD cycle at the HIV-1 reporter gene. **(a)** Sample cell showing co-localization of CTD-RNAP2, Ser5ph-RNAP2, and mRNA signals at the transcription site (TS). **(b)** The TS from (a) at select times. **(c)** Normalized intensity fluctuations at the TS for CTD-RNAP2 (red circles), Ser5ph-RNAP2 (green squares), and mRNA (blue diamonds) versus time. Times of minimal mRNA (less than 0.20 a.u.) are marked with dashed gray lines (t_1-4_). **(d)** The average normalized intensity of each signal surrounding times of minimal mRNA (40 events from 13 of 20 cells; solid connecting line shows Gaussian fit). Both the Ser5ph-RNAP2 and CTD-RNAP2 signals have deep minima well below the steady state value (dashed horizontal line). Solid vertical lines mark the minima with a lighter shadow depicting the SEM from the Gaussian fit. When the same analysis is performed at 100 random time points, no obvious minima are seen (gray diamonds). **(e)** Cropped 50-frame (50 min total) moving-average image of the TS in (a) and the fitted center position for mRNA (blue), Ser5ph-(green), and CTD-RNAP2 (red). **(f)** 50-frame moving-average XY position of each signal at the TS in (a) over time. Note the mRNA signal was used as the reference signal within the crop. **(g)** The distance between each signal in (f) over time: Ser5ph-RNAP2 to mRNA (cyan circles; (1)), CTD-RNAP2 to mRNA (Purple squares; (2)), and Ser5ph-RNAP2 to CTD-RNAP2 (orange diamonds; (3)). **(h)** The distribution of distances measured as in (g) at all TSs in all cells analyzed (sampled every 10 minutes). n=number of events/number of cells. Significance was tested using the Mann-Whitney U-test with p≤ 0.05 (*), p≤ 0.01 (**), and p≤ 0.001 (***).

Having established a well-controlled system to examine fluctuations at a single gene, we were confident in our ability to quantify the temporal ordering of RNAP2 and mRNA throughout the transcription cycle. One thing that stood out was that peaks and troughs in the mRNA signal tended to come after the peaks and troughs in the RNAP2 signals. Although there were some exceptions due to the stochasticity of the system, in some cells this behavior was seen multiple times in even single time series (for example, see valleys at t_1-4_= 16, 75, 94 and 113 min in Fig. 2b-c). To better quantify this effect, we selected all events at which the mRNA signal dropped below a threshold value, extracted all three signal channels from seven minutes before to seven minutes after each event, and aligned all signals relative to these mRNA minima event times (Fig. 2d). This analysis revealed two important aspects of the dynamics of our system. First, the analysis confirmed the signals were strongly correlated, since strong minima could be observed in all channels. These minima were significant compared to the results from unaligned signals (gray diamonds in Fig. 2d, p-values of 1.72×10^-10^ for Ser5ph- and 6.39×10^-4^ for CTD-RNAP2). Such strong correlation between mRNA production at the HIV-1 reporter and endogenous RNAP2 would suggest the reporter is not part of a larger transcriptional unit containing multiple genes. Second, the analysis indicated a temporal ordering, with both RNAP2 signals coming before mRNA by 0.96 ± 0.55 min for CTD-RNAP2 (p-value 3.65 × 10 ^−3^) and 0.88 ± 0.24 min for Ser5ph-RNAP2 (p-value 1.28 × 10^-5^). This delay makes sense because RNAP2 must escape the promoter and elongate 0.7 kb before it reaches the MS2 repeats. The CTD-RNAP2 signal also slightly preceded the Ser5ph-RNAP2 signal, although the delay was not significant at our sampling rate. This suggests nearly all RNAP2 at the locus either come in pre-phosphorylated or are rapidly phosphorylated at Serine 5 within a minute of arrival.

### Spatial organization of CTD phosphorylation at the reporter gene

RNAP2 is thought to be organized in phosphorylation-dependent clusters^7,8^. To test this hypothesis, we measured the center position in X and Y of CTD-RNAP2, Ser5ph-RNAP2, and mRNA at the reporter gene over time (Fig. 2e-g). If the hypothesis is correct, we would expect to see some spatial separation in our different RNAP2 and mRNA signals. To confirm this hypothesis, we calculated the Euclidean distance between each pair of signals. As Fig. 2g illustrates, the distances between signals changed over time, but were spatially organized such that the RNAP2 signals were significantly separated from mRNA.

Although there was considerable variation from cell to cell, this trend could be seen in the median positions from the whole population of transcription sites we tracked (Fig. 2h). Specifically, the median distance from mRNA to CTD-RNAP2 was ~181 nm compared to ~148 nm for Ser5ph-RNAP2 (p-value 0.032). Likewise, the median distance between the two forms of RNAP2 was just ~93 nm, significantly smaller than between either form of RNAP2 and mRNA (p-value<6.97× 10^-11^) (Fig. 2h and Sup. Fig. 3h). This spatial separation was consistent across the cells we analyzed (Sup. Fig. 4) and independent of the strength of transcription as gauged by CTD-RNAP2, Ser5ph-RNAP2, and mRNA signal intensities. Together these results demonstrate that RNAP2 is spatially organized within the transcription site, with active mRNA synthesis spatially distinct from clusters of CTD-RNAP2 and Ser5ph-RNAP2.

### Fluctuation dynamics and statistics are captured by a simple model of transcription bursting

We wanted to obtain a more universal picture of RNAP2 phosphorylation dynamics at the HIV-1 reporter gene. We therefore performed correlation analysis^21,34,35^ using all time points in all time series, similar to fluorescence correlation spectroscopy^36^. This technique is ideal for extracting information from noisy data provided there are a sufficient number of time series and/or time points. We began with an auto-correlation analysis, to see how long each signal remains correlated with itself given a lag time (*τ*) (Fig. 3a). The auto-correlation of each signal decays with increasing lag time and eventually flattens out near zero. We define the dwell time as the lag time at which the auto-correlation falls below 20% of its initial zero-lag value. According to this analysis, the two forms of RNAP2 had shorter average dwell times than mRNA, indicating RNAP2 was often unsuccessful in reaching the end of the gene and synthesizing an mRNA.

**Fig. 3:**
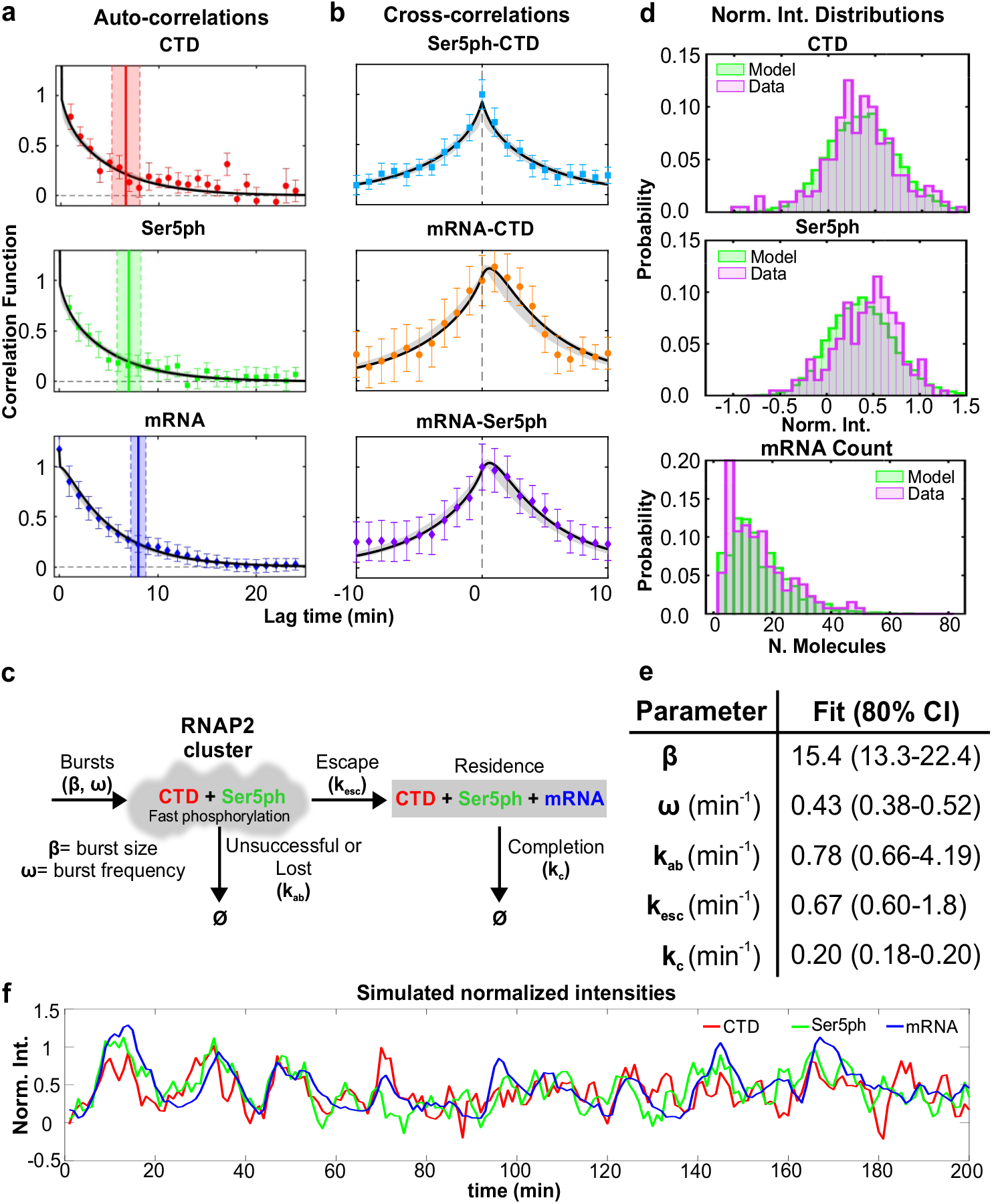
Fluorescence auto- and cross-correlations at the HIV-1 reporter gene are well fit by a unifying model of transcription. **(a,b)** Measured and modeled (**a**) auto-correlation functions 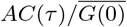 for each signal: CTD-RNAP2 (red circles), Ser5ph-RNAP2 (green squares), and mRNA (blue diamonds). Dwell time is defined as the time at which the autocovariance dropped below 20% its zero-lag value (vertical full lines). Dwell time uncertainty is estimated from the model using SD from 400 simulated data sets, each with 20 cells over 200 min with 1 min simulation resolution (vertical dashed-lines). **(b)** cross-correlation function 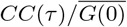 between signal pairs: Ser5ph-RNAP2 and CTD-RNAP2 (cyan squares), mRNA and CTD-RNAP2 (orange circles), and mRNA and Ser5ph-RNAP2 (purple diamonds) at the transcription site. Model MLE fit in black and uncertainty in gray. **(c)** A simple model to capture RNAP2 fluctuation dynamics at the HIV-1 reporter gene. RNAP2 enter the transcription cluster with an average geometric burst with average burst size, *β*, and burst frequency, *ω*. Phosphorylation of Serine 5 is assumed to be fast (≪ 1 min) and/or the RNAP2 enter in a pre-phosphorylated form. RNAP2 can be lost from the cluster with rate *k_ab_* or escape with rate *k_esc_.* RNAP2 completes transcription with rate k_c_. **(d)** Probability distributions for CTD-RNAP2 and Ser5ph-RNAP2 (arbitrary units of fluorescence), and mRNA (units of mature mRNA) for experimental data (purple) and model MLE predictions (green). **(e)** Maximum likelihood estimate (MLE) parameters and 80% CI range. Statistics presented for the data are sample means ± SEM. n=number of cells/number of independent experiments (20/8). **(f)** Simulated trajectory (with shot noise equal to that of experiments) of CTD-RNAP2 (red), Ser5ph-RNAP2 (green), and mRNA (blue) intensities normalized to have a 95 percentile of unity.

Next, we calculated the cross-correlation between signals. Consistent with our previous analysis aligning local minima, all possible pairs of signals were strongly correlated, as seen by large peaks in the cross-correlation curves near τ=0 (Fig. 3b). Measuring the precise position of each peak revealed the mRNA signal came substantially later than the CTD-RNAP2 and Ser5ph-RNAP2 signals, while the CTD-RNAP2 and Ser5ph-RNAP2 signals appeared at roughly the same time (within the 1 min sampling time of experiments). To better resolve the time delay between the CTD-RNAP2 and Ser5ph-RNAP2 signals, we re-imaged the HIV-1 transcription site in a single plane at a much faster frame rate (150 ms/frame) for a total of 1000 timepoints (150 sec). Although these higher temporal resolution experiments are much too short to capture the full auto- and crosscorrelation curves, they are sufficient to resolve the short time-lag dynamics (Sup. Fig. 5), and they revealed cross-correlation asymmetry with an off-center peak indicating that the Ser5ph-RNAP2 signal comes roughly 3-6 sec after the CTD-RNAP2 signal. The various delays we measure are consistent with the temporal ordering we saw by aligning local minima of the mRNA signal (Fig. 2d) and provide further evidence that RNAP2 phosphorylation at Serine 5 is very rapid at the transcription site.

We next sought to find a quantitative model to unify our diverse data sets. We required that our model must simultaneously fit all three auto-correlation curves (Fig. 3a) and all three cross-correlation curves (Fig. 3b). To further constrain the model, we also counted mRNA at transcription sites by comparing their intensities to single mature mRNAs using FISH-quant^37^ (Fig. 3d, bottom). Consistent with an earlier report^20^, we found the HIV-1 reporter contained an average of *μ* = 15.5 mRNA with a relatively large standard deviation of *σ =* 10.55, and Fano Factor of *σ*^2^/*μ* = 7.1.

To unify our data, we posed several models with different levels of complexity (Sup. Fig. 6). Each model considered a promoter with bursty expression. This was represented by specifying distinct active (ON) and inactive (OFF) promoter states with OFF-to-ON and ON-to-OFF transitions rates *k_on_* and *k_off_*, respectively. When the promoter is ON, RNAP2 is recruited at a rate *k_r_*^38^. Upon fitting these models to our data, the fitted burst duration was much shorter than the one minute experimental sampling time (i.e., *k_off_ ≪ 1min).* This allowed us to simplify the model to one with burst frequency *ω = 1/(1/k_on_ + 1/k_off_) = 1 /k_on_* and geometrically distributed bursts with average size *β = k_r_/k_off_*^39^. In all models that fit our data, RNAP2 could unsuccessfully depart the promoter at rate *k_ab_* or escape at rate *k_esc_.* After escape, the RNAP2 would complete transcription at a combined rate *k_c_* that includes both elongation and processing.

In the minimal model that matched all data, CTD-RNAP2 were immediately phosphorylated upon arrival at the promoter, which was consistent with the rapid (< 1 min) Serine 5 phosphorylation we observed (Sup. Fig. 5). We also explored several more complicated models with separate steps for initiation, elongation, and processing, post-transcriptional mRNA retention^40^, or with separate events describing Serine phosphorylation/initiation and de-phosphorylation/abortion (Sup. Fig. 6). Each model was fit separately to maximize the likelihood for all observed data, but inclusion of additional mechanisms and free parameters provided only marginal improvements to the overall fit and resulted in much larger parameter uncertainties. Therefore, we used the Bayesian Information Criteria (BIC) to select our final model as the best choice given our available data (See tables in Sup. Fig. 6). By simultaneously fitting all six correlation plots (Fig 3a and 3b) as well as the nascent mRNA means and variances, we could estimate the best model’s five parameters with excellent precision (Sup. Fig. 7). The best-fit parameter values and their uncertainties are provided in Fig. 3e. According to the best fit, bursts of RNAP2 occur on average every 1/ω ≈ 2.3 min and have an average size of about *β ≈* 15 molecules per burst. Of the RNAP2 that arrive at the promoter, a substantial fraction *f* = k_esc_/(k_esc_ + *k_ab_) ≈* 0.46 escape the promoter and complete transcription, leading to convoys^20^ of about *f* · *β* ≈ 7 RNAP2 per burst. Each mRNA takes an average of 1 */k_c_ ≈* 5 min to complete elongation and processing, meaning that on average the HIV-1 reporter contains mRNA originating from *ω/k_c_ ≈* 2 consecutive bursts. Overall, the model predicts that there are an average of ~20 RNAP2 on the gene in steady-state, with an average of ~5 in the cluster near the promoter in an unphosphorylated or Ser5ph form, and ~15 elongating or processing near the end of the gene (See Table 1). This average picture is somewhat misleading, however, as the number of RNAP2 within the cluster fluctuates dramatically due to randomly timed bursts. According to our simulations, there are periods when as many as ~90 RNAP2 come in at a time interspersed by brief and random silent periods of low RNAP2 occupancy (Sup. Fig. 8a).

**Table 1:**
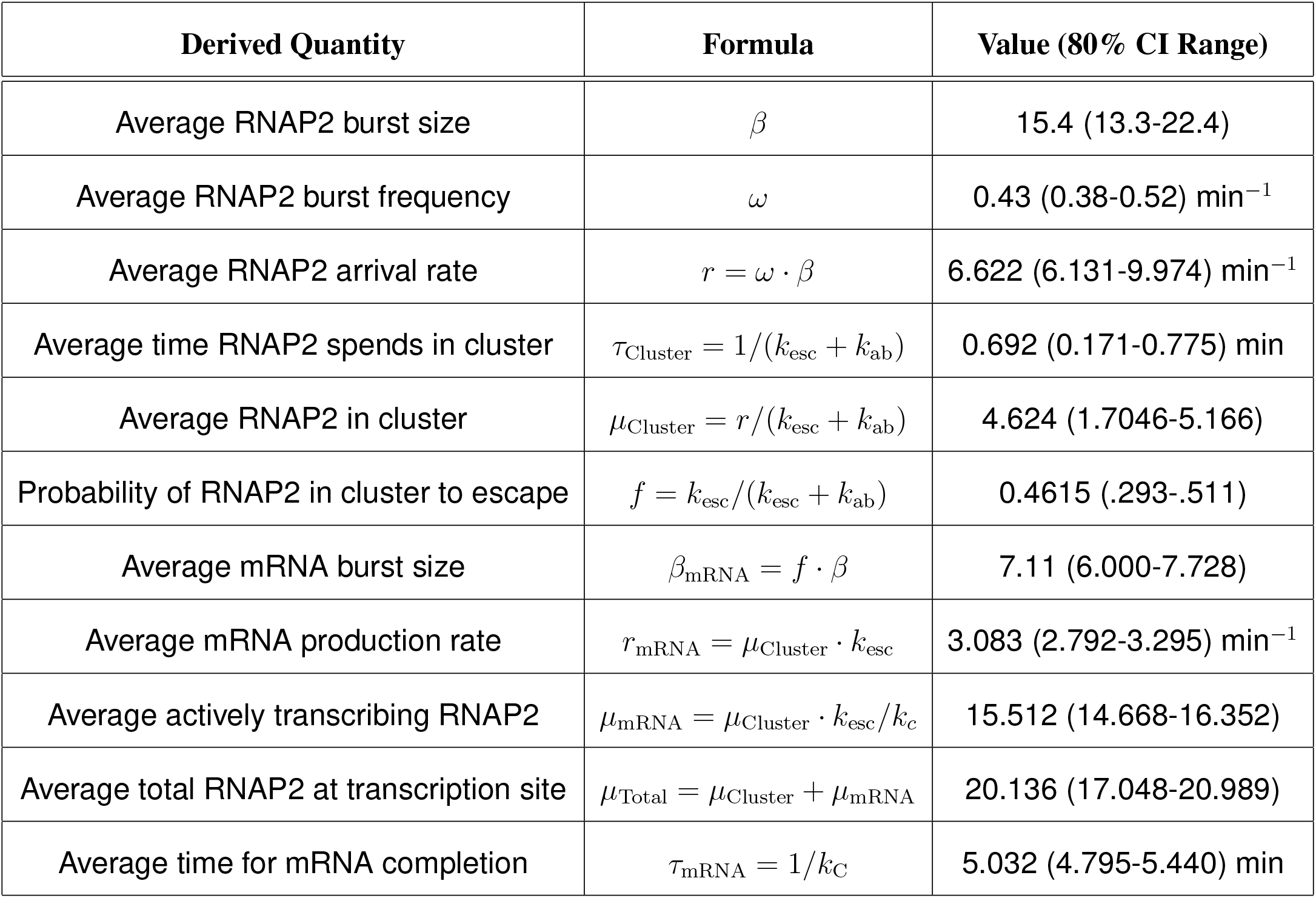
Derived Quantities and Confidence Intervals Resulting from Model Fit to Data

| Derived Quantity | Formula | Value (80% CI Range) |
| --- | --- | --- |
| Average RNAP2 burst size | $\beta$ | 15.4 (13.3-22.4) |
| Average RNAP2 burst frequency | $\omega$ | 0.43 (0.38-0.52) min<sup>-1</sup> |
| Average RNAP2 arrival rate | $r = \omega \cdot \beta$ | 6.622 (6.131-9.974) min<sup>-1</sup> |
| Average time RNAP2 spends in cluster | $\tau_{\text{Cluster}} = 1/(k_{\text{esc}} + k_{\text{ab}})$ | 0.692 (0.171-0.775) min |
| Average RNAP2 in cluster | $\mu_{\text{Cluster}} = r/(k_{\text{esc}} + k_{\text{ab}})$ | 4.624 (1.7046-5.166) |
| Probability of RNAP2 in cluster to escape | $f = k_{\text{esc}}/(k_{\text{esc}} + k_{\text{ab}})$ | 0.4615 (.293-.511) |
| Average mRNA burst size | $\beta_{\text{mRNA}} = f \cdot \beta$ | 7.11 (6.000-7.728) |
| Average mRNA production rate | $r_{\text{mRNA}} = \mu_{\text{Cluster}} \cdot k_{\text{esc}}$ | 3.083 (2.792-3.295) min<sup>-1</sup> |
| Average actively transcribing RNAP2 | $\mu_{\text{mRNA}} = \mu_{\text{Cluster}} \cdot k_{\text{esc}}/k_c$ | 15.512 (14.668-16.352) |
| Average total RNAP2 at transcription site | $\mu_{\text{Total}} = \mu_{\text{Cluster}} + \mu_{\text{mRNA}}$ | 20.136 (17.048-20.989) |
| Average time for mRNA completion | $\tau_{\text{mRNA}} = 1/k_c$ | 5.032 (4.795-5.440) min |

After fitting the model to capture the auto- and cross-correlation functions and the mean and variance of the mRNA distribution, we verified that it also correctly predicted the full probability distributions for the number of nascent mRNA molecules and RNAP2 signal intensities at the HIV-1 transcription site (Fig. 3d). We also simulated normalized intensities including shot noise (Fig. 3f), and these look similar to our measured trajectories (Fig. 2c). The shot noise was estimated directly from the experiments by comparing the observed zero-lag covariance G(0) compared to an estimate for the zero-lag autocovariance found by interpolation from the short, but non-zero time lags. These shot noise standard deviations were found to be 1.98×, 1.42×, and 0.41× of the standard deviation for CTD, Ser5ph, and MS2 signals, respectively. Finally, we simulated ChIP data for our single-gene reporter (Sup. Fig. 8b-c). To do this, we assumed an elongation rate of 4.1 kb/min (measured previously at this locus by analyzing the MS2 stochastic fluorescence fluctuations^20^) and processing rate of 0.27 min^-1^ (so elongation and processing times sum to our fitted *1/k_c_* completion time). With these rates, the CTD/Ser5ph-RNAP2 simulated ChIP signals from active genes displayed strong peaks at the beginning and end of the gene, as we observed in Fig. 1b (compare to Sup. Fig. 8b-c). Overall, the excellent match between data and simulations indicates our best-fit model faithfully captures transcription dynamics at the HIV-1 reporter. To facilitate further exploration of our model, we provide a graphical user interface (GUI) at [https://github.com/MunskyGroup/Forero_2020]. The GUI allows exploration of how each model parameter affects model predictions, including trajectories, auto- and cross-correlations, distributions of spot intensities, simulated ChIP data, and several derived quantities to describe the CTD-RNAP2, Ser5ph-RNAP2, and mRNA burst dynamics (Sup. Fig. 9).

### Inhibiting distinct steps of the transcription cycle provides further evidence for spatiotemporal organization of RNAP2 phosphorylation

So far, our collective data and modeling suggest a precise temporal ordering of transcription dynamics, beginning with the recruitment of CTD-RNAP2, followed by rapid initiation in 3-6 sec (indicated by Ser5ph-RNAP2), and promoter escape and elongation within another minute or so (indicated by mRNA). Our data also provide evidence of heterogeneity in the distribution of RNAP2 along the gene, with high concentrations near the beginning and end of the gene (Fig. 1b). To further test our system, we perturbed it by adding three different transcription inhibitors: Triptolide (TPL), THZ1, and Flavopiridol (Flav) (Fig. 4). We began by inhibiting the earliest steps in the transcription cycle to attempt to prevent the formation of the RNAP2 cluster. To achieve this we added TPL, a small-molecule inhibitor that prevents promoter DNA opening and transcription initiation by inhibiting the DNA-dependent ATPase activity of the XPB subunit of TFIIH^4,41^. TPL has also been shown to induce RNAP2 degradation on the hours timescale^42^, so we imaged for just 30 consecutive minutes to focus on the more immediate impact of TFIIH inhibition. Addition of 5 *μ*M TPL led to a rapid and dramatic loss of both mRNA and all RNAP2 signals at the transcription site within just ~ 10 min (Fig. 4a-d, and Sup. Movie 2). Consistent with our previous findings, we observed a temporal ordering in the TPL-induced run-off of RNAP2(Fig. 4c), with CTD-RNAP2 signals dropping earlier than Ser5ph-RNAP2, followed by mRNA. This ordering was observed in 7 out of 10 single cells we measured. Of these, 4 exhibited clear separation between the three traces (inset in Fig. 4c). Since steps that are later in the CTD cycle necessarily take longer to respond to drugs, this ordering provides further evidence that CTD-RNAP2 slightly precedes Ser5ph-RNAP2 by less than a minute, and that both RNAP2 signals come significantly earlier than mRNA. These data also demonstrate that the opening of promoter DNA by XPB is a requirement for the formation of RNAP2 clusters. This can work by at least two mechanisms: (1) All the Ser5ph-RNAP2 underwent initiation and abortion, but RNAP2 kept its Serine 5 phosphorylation; (2) Initiation of the first RNAP2 activates CDK7, which can phosphorylate many RNAP2 within the cluster.

**Fig. 4:**
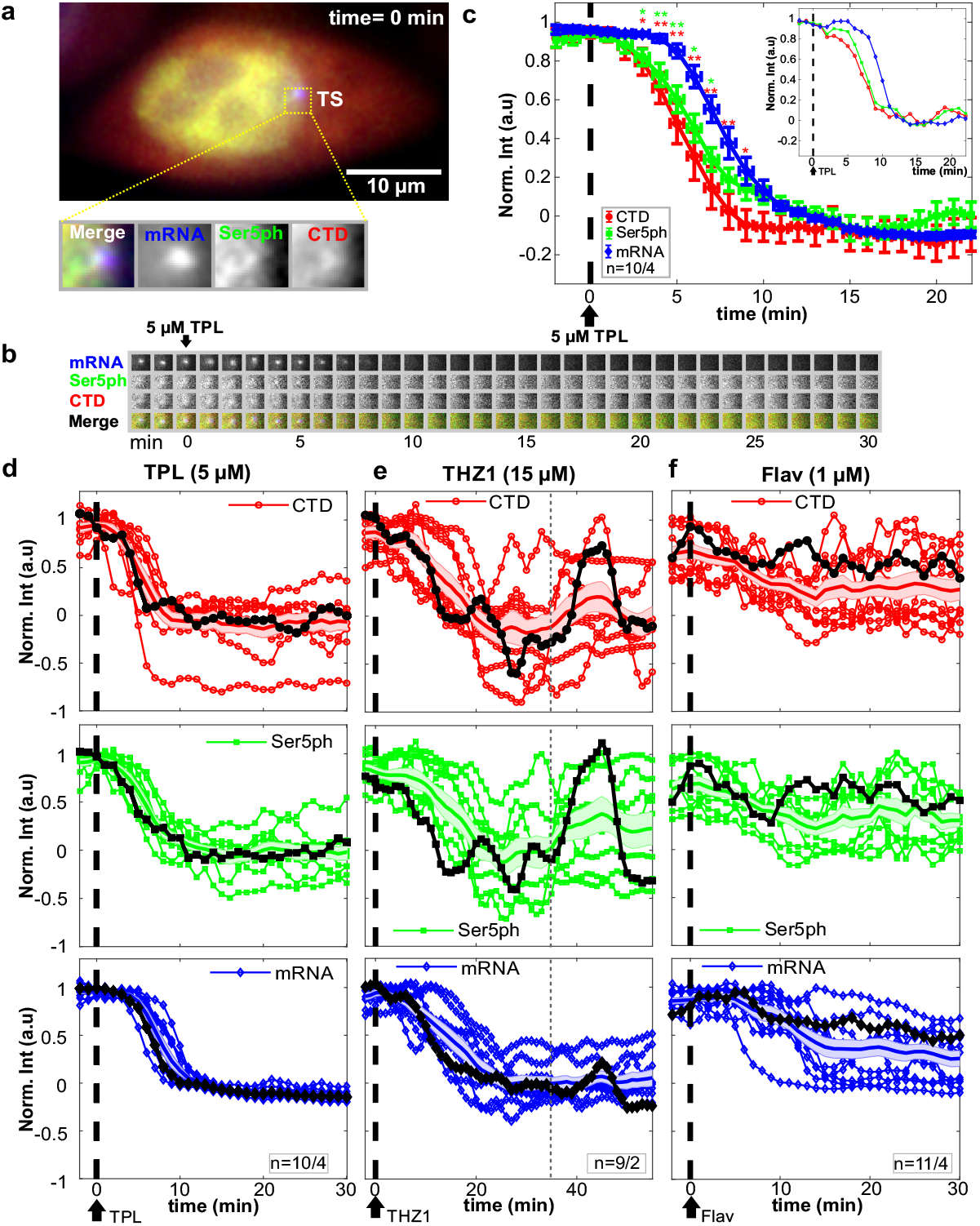
Intensity fluctuations of CTD-RNAP2, Ser5ph-RNAP2 and mRNA in the presence of transcription inhibitors. **(a)** Sample cell before addition of Triptolide (TPL). The transcription site (TS) is shown in the dotted box, and the inset shows a zoom in. **(b)** The TS from (a) at all times before and after addition of TPL. **(c)** Normalized average TS intensity over time of all the quantified cells for CTD-RNAP2 (red circles), Ser5ph-RNAP2 (green squares), and mRNA (blue diamonds) before and after application of TPL (vertical dashed line). The inset shows the signals in a representative cell. Significance was tested using the Mann-Whitney U test with p ≤ 0.05 (*), p ≤ 0.01 (**), and p ≤ 0.001 (***) noted. **(d-f)** Normalized intensity signals after application of various transcription inhibitors, including **(d)** TPL (5 *μ*M), **(e)** THZ1 (15 *μ*M), and **(f)** Flavopiridol (Flav, 1 *μ*M) for all the cells analyzed. Signals highlighted in black correspond to a sample single cell, the colored shadow and the full line in the middle of it correspond to the standard deviation and the mean in each channel, respectively. The vertical gray dashed line in (e) highlights the time point at which a burst of RNAP2 is observed in the sample cell in the presence of THZ1. n=number of cells/number of independent experiments.

We next used THZ1, which inhibits RNAP2 CTD phosphorylation at Serine 5 by targeting the TFIIH kinase CDK7, thereby preventing promoter pausing, mRNA capping, and productive elongation^4,18,43^. In contrast to TPL, THZ1 has a slower action, so a higher concentration and longer exposure to this drug were needed to see an effect in real-time. Treatment with 15 *μ*M THZ1 led to a reduction in the mRNA signal at the HIV-1 reporter within 25 min (Fig. 4e). Likewise, both CTD-RNAP2 and Ser5ph-RNAP2 levels were on average reduced. Interestingly, in some single cells we observed large, temporally ordered bursts in the levels of CTD-RNAP2 and Ser5ph-RNAP2, despite continued inhibition and overall loss of mRNA. These large bursts could even achieve RNAP2 levels that were as high as pre-treatment levels (the thicker black curve in Fig. 4e highlights one example). Presumably these bursts occur because there is residual TFIIH left in the cell that are not yet inhibited by THZ1, or because recently aborted RNAP2 retain their Ser5ph within the cluster. Since mRNA levels did not burst to the same degree, we conclude the bursts arise from clusters of RNAP2 near the promoter that initiate but fail to escape. These transient clusters near the beginning of the gene are consistent with the high concentration of RNAP2 near the promoter we observed by ChIP (Fig. 1b) and are also consistent with the ChIP predictions of our best-fit model (Sup. Fig. 8b-c).

We next blocked a later step in the transcription cycle using 1 *μ*M Flav, a drug that prevents transcription elongation and RNAP2 CTD phosphorylation at Serine 2 by inhibiting the CDK9 activity of P-TEFb^18,44^. Like THZ1, Flav also reduced the intensity of the mRNA signal, this time within ~15 min (Fig. 4f). However, CTD-RNAP2 and Ser5ph-RNAP2 signals remained relatively unchanged, exhibiting large fluctuations and a slight overall reduction on average. This difference from THZ1 can be attributed to the later action of Flav in the transcription cycle. The high levels of CTD-RNAP2 and Ser5ph-RNAP2 signals that remained post-Flav again support a dynamic clustering model^4,5,7–9^ in which most RNAP2 are already phosphorylated at Serine 5 and presumably make repeated attempts at initiation and promoter escape.

Finally, we attempted to qualitatively recapitulate these perturbations using our best-fit model. To do so, we evaluated several hypothetical mechanisms in which transcription is inhibited by reducing one or more of the rates, including burst statistics (*ω* or *β*), the promoter escape rate *k_esc_*, or the completion rate k_c_. According to simulations, inhibiting earlier steps (*ω* or *β*) in the transcription cycle led to the sequential loss of all RNAP2 and mRNA signals at the transcription site at a rate governed by the time scale of mRNA elongation and processing (Sup. Fig. 10a), reminiscent of our TPL experiments. In contrast, inhibiting a later step (k_esc_), led to a retention of large numbers of RNAP2 in the cluster that undergoes relatively large and rapidly changing fluctuations (Sup. Fig. 10b), reminiscent of our THZ1 experiments. Blocking (k_esc_) and reducing *k_c_* by 30 % led to a slight reduction in the mRNA signal and even less decrease in the RNAP2 signals with relatively large fluctuations (Sup. Fig. 10c), reminiscent of our Flav experiments. We also blocked bursts (either *ω* or *β*) and reduced *k_c_* by 30 % and obtained an overall reduction of all the signals (Sup. Fig. 10d) that do not represent any of the inhibitors tested here. The similarity between these simulations and our experimental perturbations provide further support for our model and also provide evidence that the tested inhibitors act on distinct stages of the RNAP2 transcription cycle.

## Discussion

In this study, we measured the dynamics of the RNAP2 CTD transcription cycle at the single-gene level in living cells. By combining complementary antibody-based imaging probes with multicolor single-molecule microscopy and computational modeling, we were able to detect organization in both the temporal ordering and spatial distribution of endogenous RNAP2 phosphorylation along a single HIV-1 reporter gene.

We find that a large number of RNAP2 at the HIV-1 transcription site are clustered around the promoter in a region that is spatially distinct from elongating RNAP2 and mRNA synthesis (as depicted in Fig. 5). This spatial organization supports the notion of dynamic RNAP2 clusters that form transcriptional hubs^45^ or factories^46,47^ that contain high concentrations of transcription machinery. In steady-state, we estimate there are an average of ~20 RNAP2 at the HIV-1 gene. This total number of RNAP2 is in between recent estimates of ~80 RNAP2^6^ clustered at the constitutively expressed beta-actin locus, ~17 RNAP2 at an exogenous mini-gene^17^, and ~7.5 RNAP2 at the Pou5f1 locus^17^. Of the ~20 RNAP2 at our HIV-1 reporter gene, we estimate on average ~5 are at or near the promoter, awaiting initiation or promoter escape. During frequent bursts, however, this number can dramatically increase to as high as 90 RNAP2, with most either coming in with Serine 5 phosphorylation or rapidly acquiring Serine 5 phosphorylation within seconds (Sup. Fig. 5). Given the limited amount of space at the promoter, it is hard to imagine all of these RNAP2 are promoter bound. Instead, we believe many are unbound and collectively this fraction helps form the transcription cluster, which remains spatially distinct from mRNA synthesis.

**Fig. 5:**
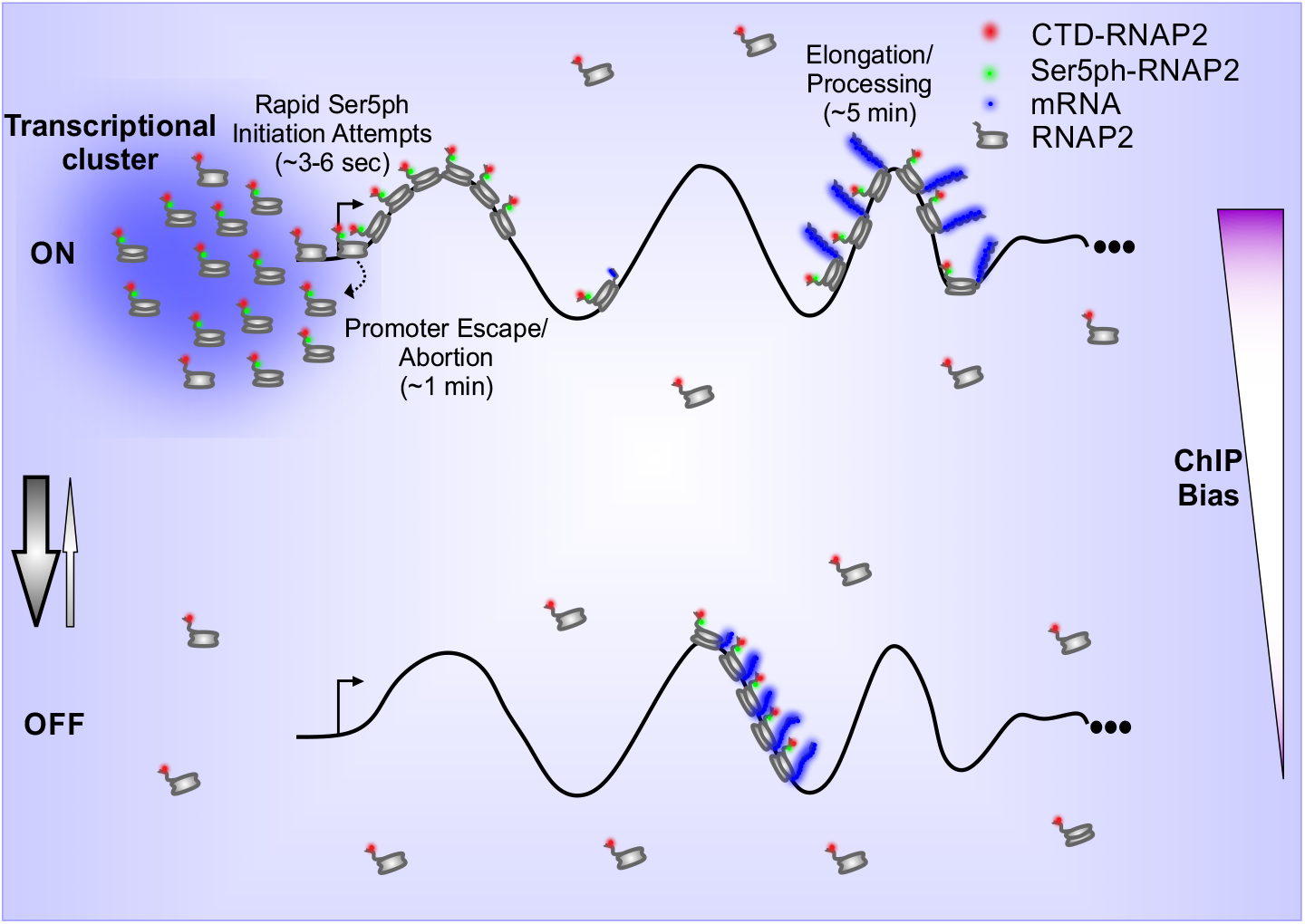
Model depicting RNAP2 transcription dynamics in a single-copy gene. During extremely short (≪ 1 min) periods of the ON state, RNAP2 is recruited in bursts (~ 15 RNAP2) the HIV-1 reporter gene, creating transient (~ 1 min) of CTD-RNAP2 and Ser5ph-RNAP2 at the gene promoter, and initiating transcription in RNAP2 convoys (~ 7 RNAP2/convoy). The middle of the gene remains mostly empty due to rapid transcription, while a large number of RNAP2 (~ 15) concentrate at the end of the gene during processing (~ 4 − 5 min). In OFF periods (~ 2.3 min), RNAP2 convoys that escaped the promoter during the ON state quickly elongate and complete transcription. The gene rapidly transitions back to the OFF state when ON (denoted by arrows). ChIP assays enrich for genes with lots of RNAP2, which will bias the assay towards genes with RNAP2 clusters near the promoter.

A major unresolved question is how RNAP2 are retained in clusters. One possibility is that RNAP2 are trapped by repeated interactions with other transcription machinery in the region. Alternatively, clusters could represent phosphorylation-dependent condensates. As others have recently shown, phase separation can be driven by phosphorylation of the unstructured RNAP2 CTD^7^ and by the histidine-rich tail of P-TEFb^8^. Since Tat directly interacts with P-TEFb^28,48^, it could enhance RNAP2 recruitment and clustering at the HIV-1 reporter gene.

One possible advantage of the cluster is it retains recently aborted RNAP2 near the transcription start site so they can rapidly re-initiate. This follows from our rapid imaging experiments, which indicate initiation is very rapid (3-6 sec; Sup. Fig. 5) compared to promoter escape (fitted *1/k_esc_ ~* 1.5 min). The distinct timescales imply two hypotheses: First, most promoter escape attempts fail. This is consistent with earlier measurements based on FRAP that demonstrated successful promoter escape is a rare event^18,49^. Second, a large fraction of RNAP2 in the cluster are inactive at any given time^5,50^. Such a large fraction of inactive RNAP2 could arise from recently aborted molecules that retain their Ser5ph. Evidence for the retention of Ser5ph on RNAP2 after transcription abortion was seen in an earlier study^18^, where Ser5ph-RNAP2 was detected in the soluble fraction of cells after transcription was globally inhibited via flavopiridol. The retention of RNAP2 also helps explain our model prediction that nearly half of the RNAP2 in the cluster *(k_esc_/(k_esc_* + *k_ab_)* ~46%) eventually do escape the promoter and produce a full-length transcript. Thus, local recycling of transcription machinery within clusters may play a role in HIV-1 biogenesis, where Tat expression provides a positive feedback loop to amplify transcription and facilitate the rapid production of viral proteins in host cells^51^.

While the overall efficiency of transcription is relatively high at the HIV-1 reporter gene compared to other genes studied, the various kinetic rates we quantified are fairly consistent with earlier work. In particular, we found RNAP2 takes around five minutes to complete transcription after promoter escape (1/*k_c_* in Fig. 3). This places an upper bound on the RNAP2 elongation and processing time. If we constrain the elongation rate to be 4.1 kb/min^20^ (~1 min for the full gene), then we can assign the remaining time (~ 4 min) to RNA processing at the 3’ end. Under these conditions, the model predicts a build up in RNAP2 at the 3’ end of the gene because processing takes longer than elongation. This buildup is consistent with our ChIP data in Fig. 1b. The estimated 4 min processing time is also consistent with an earlier estimate at this HIV-1 locus^20^, although such relatively long processing may not be representative of other genes. Similarly, the RNAP2 initiation and promoter escape rates we quantified are consistent with earlier reports, taking between a minute and a few minutes^27,49^. Finally, we also detected bursts in transcription that result in convoys of RNAP2, as previously reported^20^, and consistent with widespread bursting observed across the genome^38,52^. The global agreement between studies suggests some convergence in the field, particularly given the uniqueness of our data set, which is based on fluctuations of both MS2^21^ and RNAP2 Fab signals^27^.

The ability to image by fluorescence microscopy endogenous RNAP2 phosphorylation dynamics at a single-copy genes now makes it possible to estimate the RNAP2 distributions predicted by ChIP. ChIP studies of the RNAP2 CTD transcription cycle typically display heterogeneous distributions of RNAP2 that have distinct peaks of Ser5ph-RNAP2 near the promoter and Ser2ph-RNAP2 at the ends of genes^1,3,11^. However, based on ChIP alone, it is not clear if peaks represent the distribution of RNAP2 along single genes or instead represent a population of genes. For example, it could be that half of the genes have Ser5ph-RNAP2 paused at the beginning of the gene, while the other half have Ser2ph-RNAP2 being processed near the end of the gene. In this extreme example, no single gene would have RNAP2 at both ends. According to our best-fit model, the situation for HIV-1 is not this extreme, but the distribution of RNAP2 does depend sensitively on the timing of bursts. For example, early in a burst RNAP2 occupancy is heavily front-loaded, with all or nearly all RNAP2 at or around the promoter in a Serine 5 phosphorylated form. Since RNAP2 ChIP by design is biased towards genes with high levels of RNAP2 at the time of assay, genes that have recently burst are likely to be overrepresented in the data (Sup. Fig 5b-c). As our model demonstrates, soon after a burst, genes tend to have far more RNAP2 clustered around the promoter than the average gene (which has just five) (Fig. 5). According to this interpretation, the large Ser5ph-RNAP2 ChIP peak we observe near the promoter could arise from rapid and repeated promoter proximal initiation and/or pausing. Given the nature of ChIP, it is also possible the peak arises from RNAP2 within clusters that are non-specifically cross-linked during the fixation step. However, this latter possibility seems unlikely as promoter proximal peaks are also observed using techniques that detect and sequence nascent mRNA, such as GRO-seq, PRO-seq, and mNET-seq^53^. In the future, it will be interesting to see to what extent dynamic clustering observed in living cells correlates with promoter-proximal RNAP2 peaks observed across the genome in populations of fixed cells^54^.

Aside from HIV-1, our technology can now be used to examine RNAP2 phosphorylation dynamics at other single-copy genes. Given the high correlation between MS2 (mRNA) and RNAP2 (Fabs), in the future MS2 may not even be required. For example, by combining Fab and CARGO^55^, RNAP2 phosphorylation dynamics at any endogenous gene could be visualized without extensive genome editing. Alternatively, Fab could be combined with other labeling technologies such as lacO/lacI^56,57^, ROLEX^58^, ANCHOR^59^, or post-fixation via DNA FISH^60^ or CasFISH^61^. Beyond RNAP2, post-translational modifications to other proteins involved in transcription could also be studied in this way, including histones^23,62^. However, a few important caveats of Fab- or intrabody-based imaging should be kept in mind: First, if Fabs bind their targets with too low affinity, then there will be a large unbound fraction that will decrease signal-to-noise. For the CTD-RNAP2 and Ser5ph-RNAP2 Fab, the bound fraction was determined to be greater than 80%^27^. Second, if Fabs take too long to bind their targets, then very rapid processes can be entirely missed or their timescales will appear erroneously slow. According to FRAP, the vast majority of CTD-RNAP2 and Ser5ph-RNAP Fabs used in this study bind and rebind their targets in well under 10 sec^27^, meaning processes on the seconds time scale can be discerned, but anything shorter may be missed. Third, if Fabs are too numerous in a cell, they may compete with one another for binding and Fab targets could become saturated, both of which could interfere with the underlying biology. We introduce 1-3× 10^6^ Fab per cell^22^, far less than the ~ 1.5 × 10^7^ RNAP2 heptad repeats^27,63^. We therefore do not expect Fabs to compete or interfere. Together these three caveats place considerable constraints on experiments, but they are not prohibitive. With the continued development of Fab^23^, scFv^24^, and nanobodies^64^ for live-cell imaging, finding a suitable intrabody has become significantly easier. We therefore anticipate our technology will become a valuable new tool to study transcription dynamics at the single-gene level.

## Methods

### Cell Culture

Transcription dynamics experiments were performed in HeLa Flp-in H9 cells (H-128). The H-128 cell line generation was described previously^20^. Briefly, H-128 cells harbor an HIV-1 reporter gene tagged with an MS2X128 cassette, controlled by Tat expression. The HIV-1 reporter comprises the 5’ and 3’ long terminal repeats (LTRs) containing the viral promoter, polyA sites, as well as HIV-1 splice donor (SD1), splice acceptor (SA7) and Rev-responsive element (RRE). H-128 cells also stably express MS2 coat protein tagged with GFP (MCP-GFP), which binds to MS2 repeats when they are transcribed into mRNA. Cells were maintained in a humidified incubator at 37°C with 5% CO_2_ in Dulbecco’s modified Eagle medium (DMEM, Thermo Fisher Scientific, 11960-044) supplemented with 10% fetal bovine serum (FBS, Atlas Biologicals), 10 U/mL penicillin/streptomycin (P/S, Invitrogen), 1 mM L-glutamine (L-glut, Invitrogen) and either 400 *μ*g/mL Neomycin (Invitrogen) or 150 *μ*g/mL Hygromycin (Gold Biotechnology).

### Chromatin Immunoprecipitation and quantitative-Polymerase Chain Reaction (ChIP-qPCR)

ChIP was performed as described previously^65^ with minor modifications. H-128 cells grown in a 10 cm dish were fixed with 1% PFA in DMEM at room temperature for 5 min, neutralized in DMEM containing 200 mM glycine for 5 min and washed with PBS and NP–40 buffer (10 mM Tris–HCl, pH 8.0, 10 mM NaCl and 0.5% NP-40). Fixed cells were lysed with 360 *μ*L SDS dissolution buffer (50 mM Tris-HCl, pH 8.0, 10 mM EDTA and 1% SDS) and diluted with 1440 *μ*L ChIP dilution buffer (50 mM Tris-HCl, pH 8.0, 167 mM NaCl, 1.1% Triton 100 × and 0.11% sodium deoxycholate), supplemented with a proteinase inhibitor cocktail. After shearing chromatin using a Bioruptor UCD–200 (Diagenode) at sonications of 40 sec with 50 sec intervals, eight times at high level, the median size of fragmented DNA was 200 base pairs with a range of 50–500 base pairs. The supernatant, cleared by centrifugation at 20,000 ×g for 10 min at 4°C, was diluted with 5.4 mL ChIP dilution buffer and then incubated with 40 *μ*L sheep anti–mouse IgG magnetic beads pre-incubated with 1 *μ*g mouse anti-CTD-RNAP2 (MABI 0601), anti-Ser5ph-RNAP2 (MABI 0603) and anti-Ser2ph–RNAP2 (MABI 0602) monoclonal antibodies (Cosmo Bio USA) at 4^°^C overnight with rotation. The immune complexes were washed with low–salt RIPA buffer (50 mM Tris-HCl, pH 8.0, 1 mM EDTA, 150 mM NaCl, 0.1% SDS, 1% Triton 100 × and 0.1% sodium deoxycholate), high–salt RIPA buffer (50 mM Tris–HCl, pH 8.0, 1 mM EDTA, 500 mM NaCl, 0.1% SDS, 1% Triton 100 × and 0.1% sodium deoxycholate) and then washed twice with TE buffer (10 mM Tris-HCl, pH 8.0, and 1 mM EDTA). DNA was eluted with ChIP elution buffer (10 mM Tris-HCl, pH 8.0, 300 mM NaCl, 5 mM EDTA and 0.5% SDS). After incubation at 65°C overnight to reverse the crosslinks, DNA was purified by RNase A and proteinase K treatments and recovered using a DNA purification kit (Qiagen). For ChIP-qPCR, the immunoprecipitated DNA and total DNA were quantified by Power SYBR Green PCR Master Mix in a Mx3000P Real-Time qPCR System (Agilent Technologies). The primers used for qPCR are listed in Sup. Table 1.

### Antigen-binding fragment (Fab) generation and fluorescence conjugation

Fab preparation was performed using the same monoclonal antibodies used in ChIP experiments and the Pierce Mouse IgG1 Fab and F(ab’)2 Preparation Kit (Thermo Scientific), as described before^27^. In brief, ficin resin was equilibrated with 25mM cysteine (in HCl, pH 5.6) to digest the antibodies (CTD-RNAP2 or Ser5ph-RNAP2) into Fab. The IgG concentration used was 4 mg, and the digestion reaction was incubated for 5 h. Fab and Fc regions were separated using a Nab Protein A column (Thermo Scientific). Fabs were concentrated up to ~1 mg/mL using an Amicon Ultra 0.5 filter (10k cut-off, Millipore) and conjugated with CF640 or Cy3 (Invitrogen) dyes. For labeling Fab, 100 *μ*g of purified Fab and 10 *μ*L of 1M NaHCO_3_^-^ were mixed to a final volume of 100 *μ*L, then 2 *μ*L of CF640 or 2.66 *μ*L of Cy3 was added, and the mixture was incubated at RT for 2 h in a rotator protected from the light. The labeled Fab sample was passed through a PD-mini G-25 desalting column (GE Health care), previously equilibrated with PBS, to remove unconjugated Fab, and then the dye-conjugated Fab was concentrated up to ~1 mg/mL with an Amicon Ultra filter 0.5 (10k cut-off). The degree of labeling (DOL) was calculated using eq. 1, where *ε_IgG_* and *ε_dye_* are the extinction coefficients of IgG at 280 nm and the dye (provided by the manufacturer), *A_Fab_* and *A_dye_* are the absorbances determined at 280 and 650 or 550 nm, and CF is the correction factor for the dye at 280 nm (provided by the manufacturer). In this study, only Fabs with a DOL between 0.75 and 1 were used for live-imaging experiments.

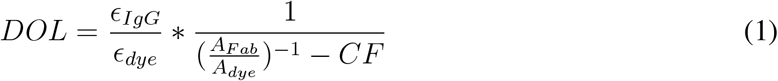

### Loading fluorescent Fabs into living cells

Cells were cultured in glass bottom dishes (35 mm, 14 mm glass, Mat-Tek). The next day dye-conjugated Fabs were loaded into the cells through bead-loading^22,27,29,66,67^, as follows: First, the fluorescent Fabs (CTD-RNAP2-CF640 and Ser5ph-RNAP2-Cy3, ~1 mg/mL, each) were mixed with PBS up to 4 *μ*L in the cell culture hood. Second, the medium was removed completely from the dish and stored, and the Fab mixture was added to the center of the dish. Third, glass beads (106 *μ*m, Sigma-Aldrich, G-4649) were immediately sprinkled on top before cells dryed up and the dish was tapped ~10 times against the bench. This tapping causes the beads to roll over cells and induce small tears into which the Fab can diffuse in. Fourth, the stored medium was quickly added back to the cells, again to prevent cells from drying out. Cells were then placed in the incubator to recover for 1-2 h. Post-recovery, the glass beads were gently washed out with phenol-free DMEM (DMEM^-^, Thermo Fisher Scientific, 31053-028), and the cells were stored in DMEM^+^ medium (DMEM^-^ supplemented with 10% FBS, 10 U/mL P/S and 1 mM L-glut) for live-imaging experiments.

### Chemicals

The transcription inhibitors, Triptolide (TPL, Sigma Aldrich), Flavopiridol (Flav, Selleck Chemicals), THZ1, (Selleck Chemicals), fluorescence dyes, Cy3 (Invitrogen), CF640 (Invitrogen), and HaloTag TMR Ligand (5 mM) (Promega) were dissolved in DMSO (Sigma-Aldrich) and stored at −20°C until use. RNAP2 inhibitors were added to DMEM^+^ medium to reach the desired final concentration in the cells.

### Microscopy

A custom-built widefield fluorescence microscope with highly inclined illumination was used in all experiments^13,33^. The microscope has three excitation beams: 488, 561, and 637 nm solid-state lasers (Vortran) that are coupled and focused on the back focal plane of the objective (60×, NA 1.48 oil immersion objective, Olympus). The emission signals were split by an imaging grade, ultra-flat dichroic mirror (T6601pxr, Chroma) and detected with two aligned EM-CCD (iXon Ultra 888, Andor) cameras by focusing with a 300 mm tube lens (generating 100× images with 130 nm/pixel). Cell chambers were mounted in a stage-top incubator (Okolab) at 37°C with 5% CO_2_ on a piezoelectric stage (PZU-2150, Applied Scientific Instrumentation). The focus was maintained with the CRISP Autofocus System (CRISP-890, Applied Scientific Instrumentation). The cameras, lasers and piezoelectric stage were synchronized with an Arduino Mega board. Image acquisition was performed with Micro-Manager software (1.4.22)^68^. Unless otherwise stated, the imaging size was set to 512 x 512 pixels^2^ (66.6 x 66.6 *μ*m^2^) and the exposure time set to 53.64 ms. The readout time of the cameras from the combination of the imaging size and the vertical shift speed was 23.36 ms, which resulted in an imaging rate of 13 Hz (77 ms per image).

For three-color imaging, far-red fluorescence (e.g. CF640 or Alexa Fluor 647) was imaged on one camera with an emission filter (FF01-731/137/25, Semrock), while red fluorescence (e.g. Cy3 or TMR) and green fluorescence (e.g. GFP) were alternately imaged on the other camera via a filter wheel (HS-625 HSFW TTL, Finger Lakes Instrumentation) with an emission filter for red fluorescence (593/46 nm BrightLine, Semrock) and green fluorescence (510/42 nm BrightLine, Semrock). The filter wheel position was rapidly switched during the 23.36 ms camera read-out time by the Arduino Mega board. For two-color imaging, far-red fluorescence was simultaneously imaged on one camera while red or green fluorescence was imaged on the other camera with the appropriate emission filters.

### Immunofluorescence

Cells grown on glass bottom dishes (35 mm, 14 mm glass, uncoated, Mat-Tek Corporation) were fixed with 4% Paraformaldehyde (Electron Microscopy Sciences) in 1 M HEPES (Sigma-Aldrich) with or without 10% Triton 100 × (Fisher Scientific) (pH 7.4) for 10 min at room temperature (RT), and washed with PBS (3×). Permeabilization (1% Triton 100 × in PBS) and blocking (100% blocking One-P, Nacalai-USA) were performed individually, for 20 min at RT, gently rocking, and rinsing with PBS (3×) after each step. The cells were incubated for 2 h at RT with 1 mL of antibody solution (10% blocking One-P:90% PBS) containing 2 *μ*g/mL of mouse monoclonal primary antibody (CTD-RNAP2 (MABI 0601), Ser5ph-RNAP2 (MABI 0603), Ser2ph-RNAP2 (MABI 0602), as described in ^27^ and now available from Cosmo Bio USA, H3K27ac (MABI0309), H3K27me (MABI0321), H3K4me1-3 (MABI0302-0304), H3K9me2 (MABI0317), and H3K9me3 (MABI0318), purchased from Cosmo Bio USA). After rinsing with PBS (3 ×), the cells were incubated for 1 h at RT with 1 mL of antibody solution containing 1.5 *μ*g/mL of Alexa Fluor 647 Donkey Anti-Mouse IgG (Jackson ImmunoResearch) and washed with PBS (3×). Then, the cells were mounted using Aqua-Poly/Mount (Fischer Scientific) for imaging. Single images were acquired with laser powers at the back focal plane set to 86 *μ*W and 51.2 *μ*W for 488 nm and 637 nm, respectively.

Additionally, to show that CTD- and Ser5ph-RNAP2 Fabs stain cells distinctively, immunostaining using pre-labeled Fabs against CTD-RNAP2-CF640 and Ser5ph-RNAP2-Cy3 was performed. For this type of experiments, cells were fixed, permeabilized and blocked as described above, and then incubated in an antibody solution containing 2 *μ*g/mL of each pre-labeled Fab for 1 h at RT. Post Fab incubation, cells were rinsed with PBS and mounted in Aqua-Poly/Mount. The images were collected using the following laser powers at the back focal plane: 123 *μ*W, 750 *μ*W, and 230 *μ*W for 488 nm, 561 nm, and 637 nm, respectively.

### Single-molecule experiments using H2B-Halo

Cells were plated in glass bottom dishes at a seeding density of ~ 10^4^ cells/cm^2^. The next day, cells were transfected with 2.5 *μ*g of H2B-Halo in a 1:1 (mass) ratio using Lipofectamine LTX (Thermofisher Scientific, 15338-100). 24 h post-transfection, cells were stained with 5nM Halo-Ligand TMR pre-treated with 30 mM NaBH_4_ for 30 min in the CO_2_ incubator (Acros Organics) to reduce the fluorophore and induce stochastic photoblinking in live-cells^69^. After staining, the cells were washed 3 times total. Each wash consisted of 3× 1mL DMEM^-^, and 1mL DMEM^+^ with a 5 minute interval between washes. Cells were imaged immediately after staining and washing. For this, the imaging size was set to 256 x 256 pixels^2^ (33.3 x 33 3 *μ*m^2^) and the exposure time set to 30 ms. This resulted in an imaging rate of 22.8 Hz (30 ms exposure + 13.86 camera readout = 43.86 ms per frame). Single z-planes were acquired for 10,000 frames total with laser powers at the objective’s back focal plane set to 125 *μ*W, and 9.93 mW for 488 nm and 561 nm, respectively. To minimize photobleaching, the 488 nm laser fired once every ten frames (to track the transcription site), while the 561 nm laser fired every frame (for tracking individual H2B).

Single molecule tracks were identified using TrackMate 3.8 with the following parameters: LoG Detector; Estimated Blob Diameter: 5.0; Pixel Threshold: 100; Sub-Pixel Localization: Enabled; Simple LAP Tracker; Linking Max Distance: 3 pixels; Gap-Closing Max Distance: 2 pixels, Gap-closing Max Frame Gap: 1 frame. Custom Mathematica code was used to calculate average Euclidean displacement for each track longer than 5 frames. Tracks were plotted with a blue-purple color distribution based upon their average Euclidean displacement. The transcription site was identified using TrackMate, and plotted in red.

### Live-cell imaging of transcription at the HIV-1 reporter gene

To cover the entire cell nucleus, all movies were taken using 13 z-stacks with 0.5 *μ*m spacing. The z position was moved only after all three colors were imaged in each plane. This resulted in a total cellular imaging rate of 0.5 Hz (2 s per volume). Note that the color scheme of the signals described in the text and figures is based on the color of the excitation lasers, CTD-RNAP2 in red (CF640), Ser5ph-RNAP2 in green (Cy3), and mRNA in blue (GFP). For shorter live-cell imaging as in Fig. 1, each cell was scanned every 1 min for 30 min with the laser power at the objective’s back focal plane set to 21.4 *μ*W, 60.5*μ*W, and 21.74*μ*W for 488 nm, 561 nm, and 637 nm, respectively, and the exposure time was 53.64 ms. For longer live-cell imaging as in Fig. 2, cells were imaged every 1 min for 200-time points, using weaker laser powers (1.15 *μ*W, 15.7 *μ*W, and 5.2 *μ*W for 488 nm, 561 nm, and 637 nm, respectively) and longer exposure times (200 ms exposure). For faster live-cell imaging as in Sup. Fig. 5, each cell was scanned at a much faster frame rate (150 ms/frame) for a total of 1000 time points (150 sec) in a single plane. For this, the imaging size was set to 256 x 256 pixels^2^ (33.3 x 33.3 *μ*m^2^) and the exposure time set to 53.64 ms, with the laser power at the objective’s back focal plane set to 1.2 mW, 335 *μ*W, and 77.5 *μ*W for 488 nm, 561 nm, and 637 nm, respectively.

### Calibrating the number of mRNA per transcription site

To count the number of nascent mRNAs at the transcription site, cells were imaged for a single time point using a higher laser power for 488 nm (230 *μ*W at the back focal plane) and a lower camera gain. These conditions allowed us to visualize both a single transcription site and single mature mRNAs. To calculate the number of mRNA per transcription site (see Fig. 3d, bottom panel): (1) Several cells were imaged on independent days. To avoid bias, cells were chosen with the same imaging conditions used for longer live-cell experiments; (2) Images were analyzed using FISH-quant V3^37^. Mature mRNAs were detected, localized in 3D with a Gaussian fit, and then a point-spread function was applied to discard spots that were larger than diffraction-limited spots. An image showing the average intensity of the mature mRNAs was created and compared to that of the transcription site. This ratio of these gave the number of nascent mRNA at each transcription site, from which the distribution shown in Fig. 3d (bottom panel, purple distribution) was computed.

### Quantifying signal intensities at the transcription site from live-cell imaging movies

Images were pre-processed using either Fiji^70^ or custom-written batch processing Mathematica code (Wolfram Research 11.1.1) to create 2D maximum intensity projections from 3D movies. Using Mathematica code, the 3D images were corrected for photobleaching and laser fluctuations, z-stack by z-stack, by dividing the movie by the mean intensity of the whole cell or the nucleus in each channel. The offset between the two cameras was registered using a built-in Mathematica routine FindGeometricTransform, which finds a transformation function that aligned the best fitted positions of 100 nm diameter Tetraspeck beads evenly distributed across the image field of view. 2D maximum projections and 3D image sequences from the images corrected for bleaching and laser fluctuations were then analyzed with a custom-written code in Mathematica to detect and track the transcription site. Briefly, thresholds were selected in each channel to visualize spots at the transcription site and a bandpass filter was used to highlight just the transcription site in the mRNA channel. The resulting image was binarized and used to create two masks for each time point: one marking the transcription site (transcription site mask: a mask semi-manually thresholded to cover just the transcription site within the image) and one marking the background (BG mask: a ring of width one pixel that surrounds the transcription site and is separated from the site by two pixels). The built-in Mathematica routine ComponentMeasurements-IntensityCentroid was used to find the coordinates of the transcription site in XY through time. The Z coordinate was determined by selecting the z-stack at which the particle in the XY coordinate had its maximum brightness (“best z”). If the transcription site disappeared (due to transcription turning off or inhibition), the Z was replaced by the Z coordinate of the last visible position. From the XYZ coordinates at each time point, a new 2D maximum projection was created considering the “best z” at each time point. From this, the pixel intensity values were recorded for each transcription site (TS) and background (BG) mask, representing the mean intensity values over time at the transcription site and the background, respectively. The raw and normalized intensity vectors were calculated per channel and a moving average of three time points was used to display the intensity *RawInt_Ch_* as a function of time, as shown in Eq. 2:

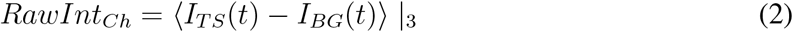

The normalized intensity (as in Fig. 2c) was calculated by dividing RawInt*ch* by the average 95% intensity from all transcription sites. Occasionally, normalized intensities for CTD-RNAP2 and Ser5ph-RNAP2 dip below zero. This can be caused either by RNAP2 signals being temporally depleted at the transcription site relative to the background or by bright signals in the background due to nearby transcription in the local vicinity. To display transcription sites over time (as in Figs. 1c and 2b and Sup. Movie 1), 3 time point moving-average trims from the “best z” were created in each channel (showing CTD-RNAP2, Ser5ph-RNAP2, mRNA, and the merge). Each trim was centered on the intensity centroid of the mRNA.

### Covariance analysis in Sup. Figs. 1f and 2g

To test for covariance between intensity signals from control spots and the transcription site, signal covariance was calculated using the “cov” function in MATLAB. For quantification of bleed through, the covariance was calculated between all possible pairs of raw intensities (CTD-mRNA, CTD-Ser5ph, and Ser5ph-mRNA) in normal vs. bleed-through control conditions. For quantification of signals off-target, the covariance was calculated between all possible pairs of normalized intensities on-target at the transcription site vs. off-target at a random site p1. Significance was calculated using the Mann-Whitney U-test.

### Analysis of minima signal in Fig 2d

The local minima in the mRNA signal of each cell was detected using the ‘islocalmin’ function in MATLAB. The cells that exhibited minimas below a threshold (normalized intensity ≤ 0.20 a.u.) were selected by the algorithm. Then 7-time points before and after the mRNA valley were considered, including the minimum, in each channel. All the traces in each channel were averaged and fitted with a Gaussian using a 95% confidence interval to determine the minima and maximum steady state of the average trace in each channel.

To confirm that the minima were true and not an artifact of our analysis, the analysis was repeated at hundreds of random time points. Significance was calculated using the Mann Whitney U-test. The p-values for the magnitude of the minima and their time delays were calculated by comparing the magnitude of the minima to the control and the time lag to minute zero in each signal, respectively.

### Analysis of transcription site spatial organization in Figs. 2e-h and Sup. Fig. 3h

Moving average (50 time points) movies were generated to accurately determine the mean XY position of the transcription site in each channel. As described in *Quantifying signal intensities at the transcription site from live-cell imaging movies* above, the built-in Mathematica routine ‘ComponentMeasurements-IntensityCentroid’ was used. Once the XY positions for each signal were obtained, the Euclidean distance between each pair over time was calculated, from which distributions were calculated. Significance between signals was calculated using the Mann-Whitney U-test.

### Auto- and cross-correlation analysis

The auto- and cross-correlation functions were calculated for each time trace obtained from the longer movies (like in Fig. 2c, but without performing a 3-time point moving-average), as previously described^34,35^. The covariance function is defined:

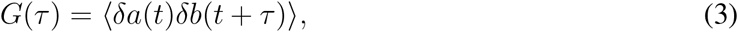

where 〈·〉 indicates the temporal mean, and *δα(t)* denotes the deviation about the mean, i.e., *δα(t) = (a(t) – 〈a(t)〉).* Signals *a(t)* and *b(t)* can be the same signal or two different signals. In the first case, *a = b* and *G(τ)* represent the auto-covariance, which is symmetric about *τ* = 0; In the second case, G(τ) represents the cross-covariance and may be asymmetric.

To calculate the cross-correlation between the CTD-RNAP2 and Ser5ph-RNAP2 signals in the fast imaging experiments in Sup. Fig. 5, the intensities of tracked transcription sites through time were quantified as described above, with a couple of minor modifications: First, because imaging was in a single plane, the rate of photobleaching in the plane was not captured by the rate of photobleaching in the cell. For this reason, each signal exponentially decayed. This was corrected by dividing out a single-exponential fit to each curve. Second, we did not perform any moving average on the signals to maintain the highest possible temporal resolution.

For fitting and data analysis, the normalized covariances, 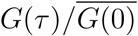, were used for all signals, where 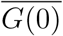 denotes the zero lag auto- or cross-covariance averaged over all time points and all biological replicas. To quantify and remove shot noise from the zero-lag auto-covariances, G(0) was estimated for each biological replica assuming a linear interpolation from the three shortest non-zero lag times (1, 2, 3 minutes) prior to averaging over all replicas. The standard error of the mean normalized covariance functions, denoted 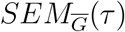, was computed as the standard deviation of 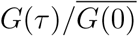 divided by 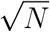.

### A quantitative model for transcription

The derivation of the bursting model for RNAP2 recruitment and nascent transcription simple model begins with the specification of three variables: *x_1_t)* describes the promoter state, *x_2_(t)* describes the number of RNAP2 in the cluster, and *x_3_(t)* describes the number of RNAP2 engaged in active transcription. Six reactions can occur: (1) a promoter can become temporarily active with propensity equal to the burst frequency, *ω* ~ *k_on_*; (2) the active promoter can deactivate at a rate *k_off_*; (3) the active promoter can recruit and phosphorylate RNAP2 at Serine 5 (Ser5ph-RNAP2) at a rate *β* · *k_off_*; (4) Ser5ph-RNAP2 can be lost from the cluster at rate *k_ab_*; (5) Ser5ph-RNAP2 can escape at rate *k_esc_*; and (6) escaped RNAP2 can complete transcription with rate *k_c_*. We solve the model for the first and second order statistical moments as previously described^71^. First, we combine the stoichiometry vectors for all six reactions into the stoichiometry matrix, S as follows:

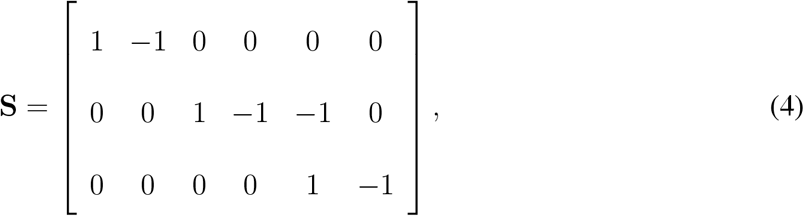

and we write the linear propensity functions in vector from as:

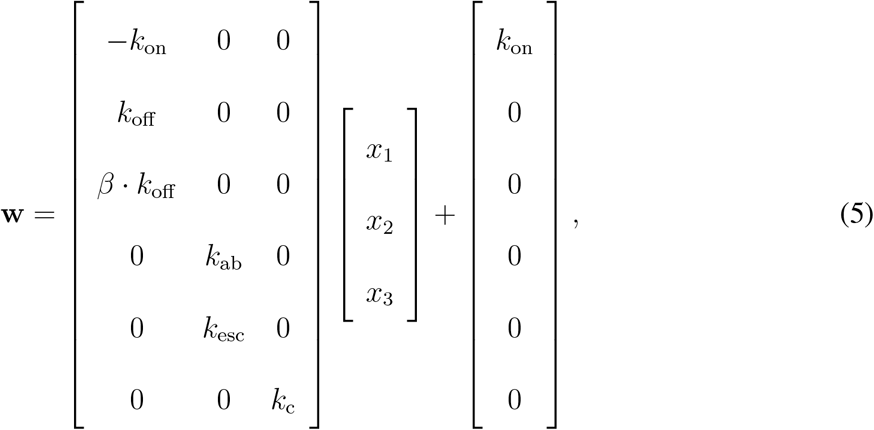

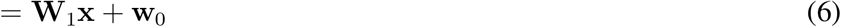

With this notation, the expected mean dynamics of 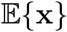 are described by the ordinary differential equation:

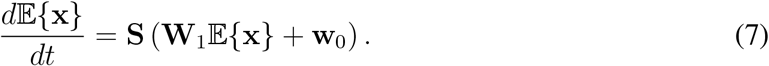

From this expression, the steady state expected mean can be calculated as the solution to the algebraic expression:

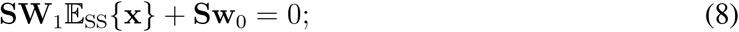

the steady state co-variance, ∑_ss_, can be calculated as the solution of the algebraic Lyapunov equation:

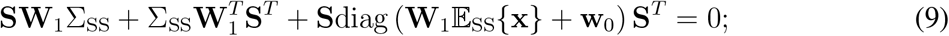

and the auto- and cross-covariance functions versus time lag, Σ(τ) can be calculated as the solution of the ODE:

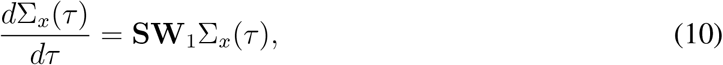

with initial condition *Σ_x_* (0) = Σ_ss_ given as the solution to (9).

To convert these above expressions, which are in terms of *x_1_, x_2_*, and *x*_3_, into quantities reflecting the total RNAP2 at the transcription site (*y*_1_ = *x*_2_ + *x*_3_) and number of transcribing RNAP2 (*y*_2_ = *x*_3_), we define a simple linear transformation:

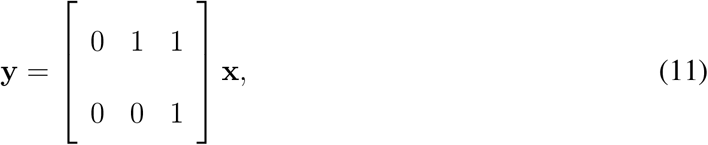

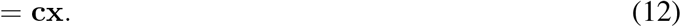

Under this transformation, 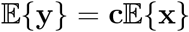 and Σ_y_(τ) = *c*Σ_x_(τ)c^T^.

We note that this version of the model does not distinguish between RNAP2 and Ser5ph-RNAP2. These two distinct forms as well as other configurations are easily incorporated by extending x to include a fourth or more states. In such cases, each new state adds two reaction stoichiometry vectors to Eqn. (4), two reaction terms to Eqn (5), and one additional row to the output matrix c in Eqn (11), but the rest of the analysis remains unchanged.

Using this model formulation, it is straightforward to solve for the steady state moments (Eqs. 8 and 9) and the auto- and cross-correlations (Eq. 10) for any combination of parameters. However, upon fitting this model to the data, we observed that estimates for *k_off_* tended to very large values (k_off_ ≫ *ω*) and with substantial estimation uncertainty. Under these excessively large rates for *k_off_*, each ‘on’ period is extremely short-lived and attracts a geometric number of RNAP2 with mean *β* (this model reduction is equivalent to the strategy in models that use geometric bursts of protein to replace translation of short-lived mRNA as described previously elsewhere^39^). Therefore, to reduce the number of free parameters required by the model, we fixed *k_off_* at 1000 min^-1^ such that each burst would be very short lived on the time scale of the experimental measurements. This choice led to simpler model, but had no discernible effect on the fit of the model to the data.

All codes, including graphical user interface will be made available on GitHub upon acceptance of the manuscript at https://github.com/MunskyGroup/Forero_2020. Advance versions are available upon request for review purposes.

### Model Parameter search

Parameters were found using maximum likelihood estimation (MLE) considering several data types as follows. First, errors in the measurement of the normalized auto- and cross-covariances were assumed to be normally distributed with the measured standard error, 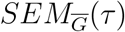, such that their log-likelihood functions are written

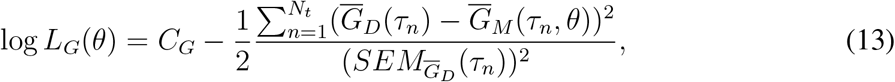

where *θ* is the set of parameters, 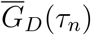 is the measured covariance function in the data (*D*), 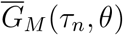 is the predicted covariance function of the model (*M*) at a time lag of *τ_n_*, and *C_g_* is a normalization constant that does not depend on the parameters. The summation is over the first 15 lag times for the three auto-covariance functions and the 21 smallest lag times (i.e., −10min to 10min) for the three cross-covariance functions.

The model was further constrained to match the mean and variance for the measured number of mRNA per transcription site as estimated in units of mature mRNA as calibrated using FISHquant. Assuming the central limit theorem, the log-likelihood of matching the observed sample mean was estimated as:

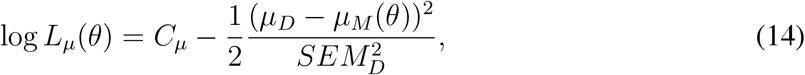

where *μ_D_* is the sample mean levels of mRNA from the data, *μ_M_* (θ) is the mean number of mRNA predicted by the model, and *SEM_D_* = 0.93 is the standard error of mean level of mRNA from the data. Similarly, the log-likelihood of the measured variance, 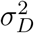 given the model was estimated as

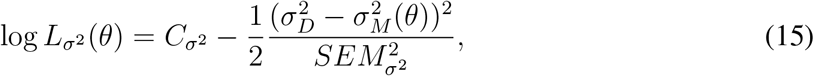

where 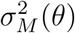 is the mRNA variance predicted by the model, *SEM_σ^2^_* is standard error for the mRNA variance, and *C_σ^2^_* is a constant that does not depend on the parameters. The standard error of the sample variance was estimated using a Gaussian approximation such that:

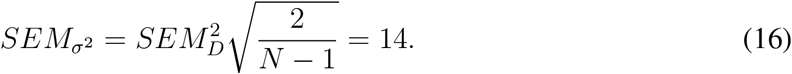

Under the assumption of independence between the different data types, the total log likelihood to match all data was the sum of the individual likelihoods:

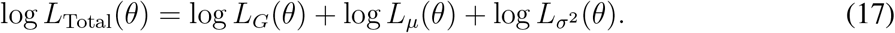

Maximum likelihood estimates were found using iterated rounds of MATLAB’s fminsearch until convergence.

To compare multiple models with different numbers of mechanisms and parameters, we computed the Bayesian Information Criteria (BIC) as:

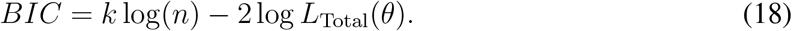

In this formulation, the value for the number of independent experiments, *n*, was estimated at *n* = 8, which conservatively assumes one data degree of freedom for each of the six different auto- and cross-correlation signals estimated from the time lapse experiments, and one each for the measurement of the mean and variance of mRNA per transcription site as estimated imaging a single frame using higher laser power to visualize single mature mRNAs. The number of parameters, *k,* disregards the directly measured shot noise magnitudes and any parameter that was fixed at a large value (e.g., *k_out_* when that values was fixed to 1000 s^-1^). This leaves *k* = 5 parameters for the selected model: *β*, *ω*, *k_ab_, k_esc_,* and *k_c_.* The Fractional Phosphorylation or Phosphorylation models each have one additional parameter, *fraction* or *k_phos_,* respectively. The mRNA retention model has one additional parameter (i.e., *k_c_* was replaced with *k_c-mRNA_*, and *k_release_* was added). The numbers of parameters, maximum likelihood values and parameter estimates, and BIC results for all examined models are listed in Sup. Fig. 6. We note that our low conservative estimate for n = 6 (rather than basing n on the much larger number of independent experiments) was chosen to avoid biasing the model selection toward simpler models - larger choices of *n* would result in much stronger rejection of the more complex models.

### Transcription inhibition experiments

For the transcription inhibition experiments in Fig. 4, cells were imaged every 1 min for 5 time points before applying the inhibitor (t=0), TPL (5 *μ*M), THZ1 (15 *μ*M), or Flav (1 *μ*M). Cells were then imaged every 1 min for 30 min total after addition of TPL or Flav, and for 55 min total after addition of THZ1. Here, laser power at the objective’s back focal plane were set to 21.4 *μ*W, 60.5μW, and 21.74μW for 488 nm, 561 nm, and 637 nm, respectively, and the exposure time was 53.64 ms.

To quantify time delays in the TPL-runoff assay, TPL signals were further analyzed as follows: (1) To account for cell variability and experimental conditions, the decays curves from each cell were aligned. This was achieved by subtracting the time at which each cell reached half of the decay after TPL addition. This time was obtained by an inverse hyperbolic tangent fit applied to each channel in every cell (Fig. 4c); (2) After the alignment, all the traces in each channel were averaged together, and the standard error of the mean (S.E.M) was calculated. Finally, to determine the time delays between CTD-RNAP2, Ser5ph-RNAP2, and mRNA, an inverse tanh fit was applied, and weighted with respect to the variance of each signal.

## Acknowledgements

We thank all the members of the Stasevich and Munsky labs for helpful discussion and suggestions, especially Luis Aguilera for help with the spot minima analyses. TJS, TM, and MS were supported by an award to TJS from the NIH (R35GM119728). BM and WR were supported by an award to BM from the NIH (R35GM124747). LSFQ was supported by the W.M. Keck Foundation. HK and TH were supported by an award to HK from JSPS KAKENHI JP17H01417 and JP18H05527.

## Author Contributions

Conceptualization, LSFQ, BM, and TJS; Performed experiments/collected data, LSFQ, TH, and MS; antibodies, HK and TH; Fab preparation, TJS, LSFQ, and HK; H-128 cells, EB; Software implementation and development, LSFQ, WR, TM, BM, and TJS; Formal analysis, LSFQ; Computational modeling, WR and BM; Wrote the original draft, LSFQ, WR, BM, and TJS; Review and edit drafts, LSFQ, WR, TH, MS, TM, EB, HK, BM, and TJS; Resources, Supervision and Funding acquisition, BM and TJS.

## Supplementary Information

**Sup. Fig. 1:**
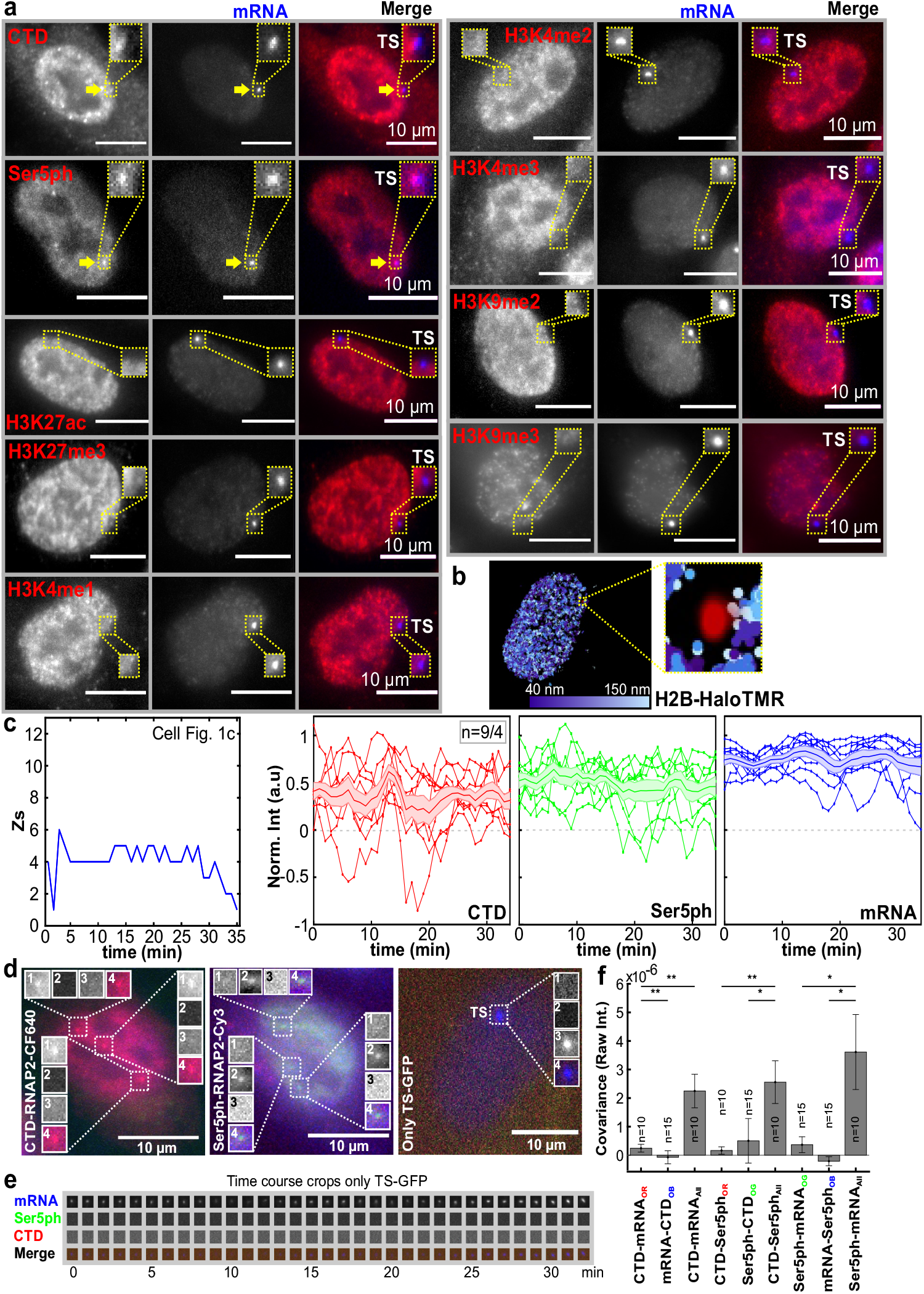
Immunostaining, single H2B tracking, and control experiments for photobleaching and bleed through. **(a)** Immunostaining (red, left panels) of CTD-RNAP2, Ser5ph-RNAP2, histone H3K27ac, H3K27me3, H3K4me1-3 and H3K9me2-3 at the HIV-1 transcription site (TS) marked by MCP-GFP (mRNA; blue, center panels), along with a merge (right panels). **(b)** Representative cell showing a mobility map of single H2B tracks. The blue scale shows the average frame-to-frame jump size (one frame every 43.86 ms) for each tracked molecule. The track corresponding to the transcription site is shown in red. The yellow dashed box displays a zoom-in around the transcription site region, where H2B is depleted. Control experiments for photo-bleaching showing **(c)** left panel, “best-Z” positions of the TS over time for the exemplary cell in Fig. 1c; right panels, normalized intensity over time for CTD-RNAP2 (red circles), Ser5ph-RNAP2 (green squares), and mRNA (blue diamonds) for all the cells recorded as in Fig. 1c,d. The shadow and the line in the middle represent the S.E.M and the average. **(d)** Images of cells from bleed-through control experiments. Left, a cell loaded with just Fab marking CTD-RNAP2 (CTD-RNAP2-CF640) displays endogenous puncta that are not the TS (designated Only Red “OR” spots); Middle, a cell loaded with just Fab marking Ser5ph-RNAP2 (Ser5ph-RNAP2-Cy3) displays endogenous puncta that are not the TS (designated Only-Green “OG” spots); Right, a cell without Fab in which the TS is marked solely by GFP-MCP binding mRNA (Only TS-GFP; Only Blue “OB”). Cropped images show the various “OR”, “OG”, and “OB” sites where the individual channels are separated and labeled as follows: (1) Red (CTD-RNAP2), (2) Green (Ser5ph-RNAP2), (3) Blue (mRNA), and (4) Merge. **(e)** Cropped images in a time course at an “OB” site demonstrates no bleed through of the mRNA channel into the other channels. **(f)** Covariance between all possible pairs of raw intensity signals is not significant at “OR”, “OG”, and “OB” sites, but is significant at the TS in cells containing all three signals (i.e. cells loaded with both Fab and expressing MCP-GFP; All). n=number of cells/number of independent experiments. Significance was tested using the Mann-Whitney U-test and noted as p-values, p≤ 0.05 (*), p≤ 0.01 (**), and p≤ 0.001 (***).

**Sup. Fig. 2:**
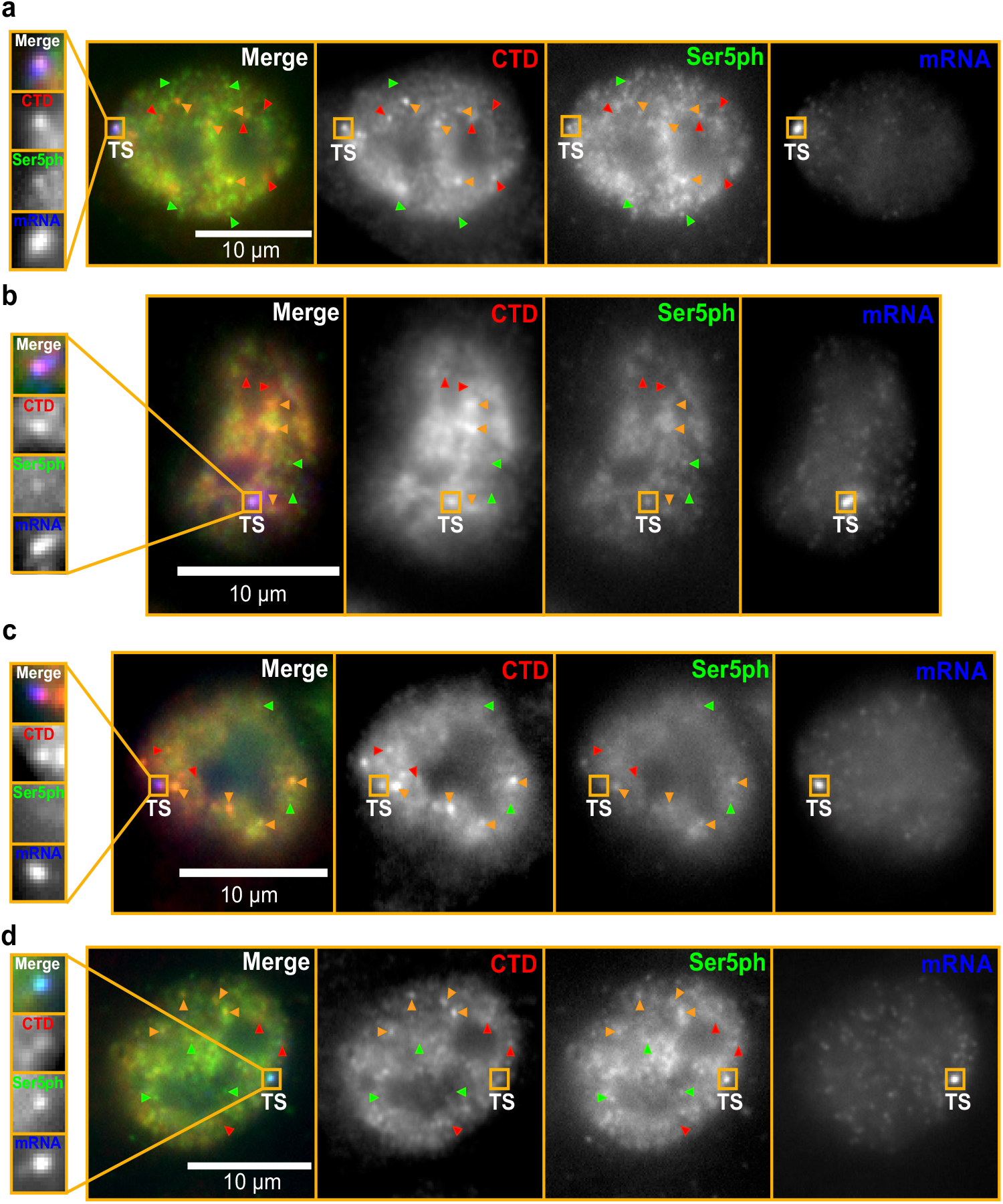
Fixed cells stained with our CTD- and Ser5ph-specific Fab. Cells with distinct staining patterns. Some areas within cell nuclei are enriched with CTD-specific Fab (red arrows), other areas are enriched with Ser5ph-specific Fab (green arrows), while still other areas are enriched with both Fabs (orange arrows). At the HIV-1 transcription site (TS), we typically see both Fabs present **(a)**. However, on occasion we can find TSs in which Ser5ph-RNAP2 staining is relatively dim **(b & c)** or, in very rare cases, CTD-RNAP2 staining is relatively dim **(d)**. This provides evidence that the signals are fluctuating.

**Sup. Fig. 3:**
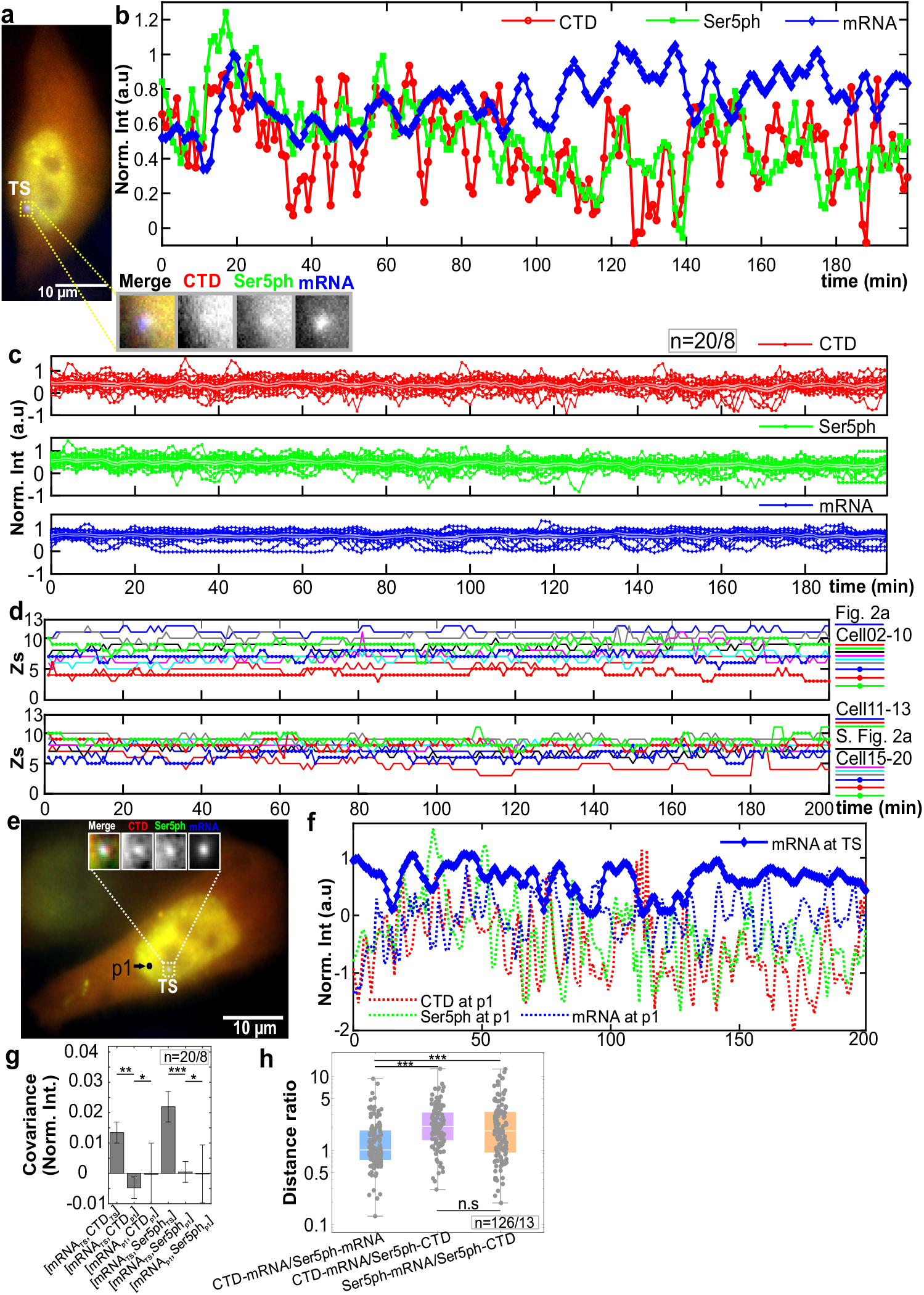
RNAP2 fluctuations at the HIV-1 reporter locus and off target. **(a,b)** A cell with strong and persistent transcription has co-localized CTD-RNAP2 (red circles), Ser5ph-RNAP2 (green squares), and mRNA (blue diamonds) at the transcription site (TS). **(c)** Normalized signal intensities over time for all the cells analyzed as in b. The shadow and the line in the middle represent the S.E.M and the average, respectively. **(d)** Z-positions of all the cells quantified for transcription fluctuations. Each cell is represented with a different color/symbol (legend on the right). **(e)** Exemplary cell with periods of active and inactive transcription showing a control position (p1) near the transcription site. **(f)** Normalized intensity over time-target at an off-target position near the transcription site (p1; CTD-RNAP2, dashed red; Ser5ph-RNAP2 dashed green; mRNA, dashed blue) versus the mRNA signal at the transcription site (blue diamonds). **(g)** Covariance calculation between the normalized intensities of mRNA at the transcription site against CTD-RNAP2 or Ser5ph-RNAP2 at the transcription site and at p1. **(h)** Ratiometric distribution of the euclidean distances for CTD-RNAP2 and mRNA to Ser5ph-RNAP2 and mRNA (light blue), CTD-RNAP2 and mRNA to Ser5ph-RNAP2 and CTD-RNAP2 (light purple), and Ser5ph-RNAP2 and mRNA to Ser5ph-RNAP2 and CTD-RNAP2 (light orange) in all the cells analyzed. Statistics values are presented as means ± S.E.M. n=number of cells, events/number of independent experiments. Significance was tested using the Mann-Whitney U-test noted as p≤ 0.05 (*), p≤ 0.01 (**), and p≤ 0.001 (***).

**Sup. Fig. 4:**
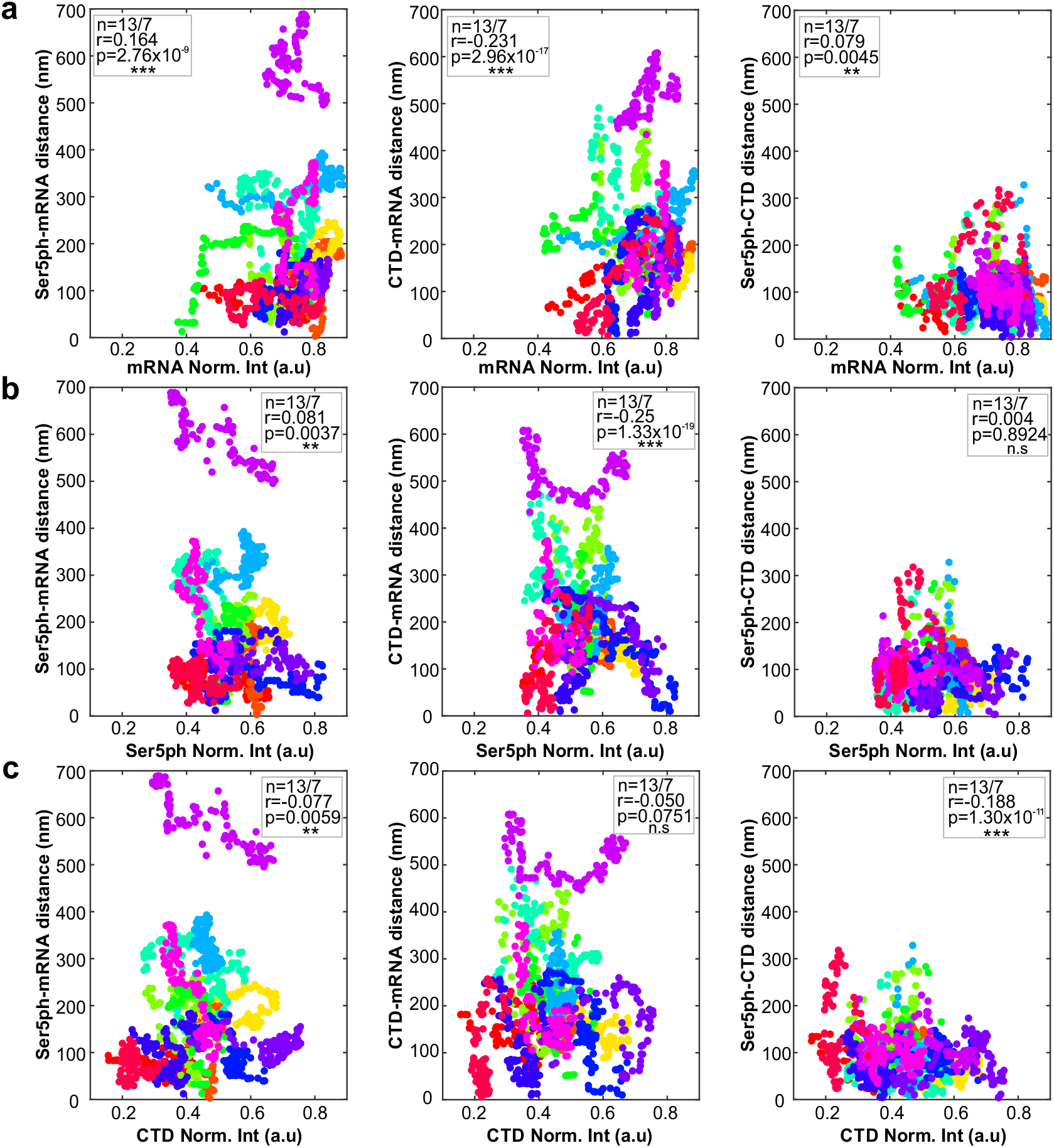
Euclidean distances distribution versus mRNA, Ser5ph-RNAP2, and CTD-RNAP2 normalized intensities. Euclidean distance between Ser5ph-RNAP2 and mRNA (left panel), CTD-RNAP2 and mRNA (middle panel), and Ser5ph-RNAP2 and CTD-RNAP2 (right panel) versus the normalized intensities of **(a)** mRNA, **(b)** Ser5ph-RNAP2, and **(c)** CTD-RNAP2 for all the cells analyzed. Each cell corresponds to one color. n=number of cells/number of independent experiments. Correlation coefficient (r) and p-values (p) as, p≤ 0.05 (*), p≤ 0.01 (**), and p≤ 0.001 (***) were calculated using the “corrcoef” function in MATLAB.

**Sup. Fig. 5:**
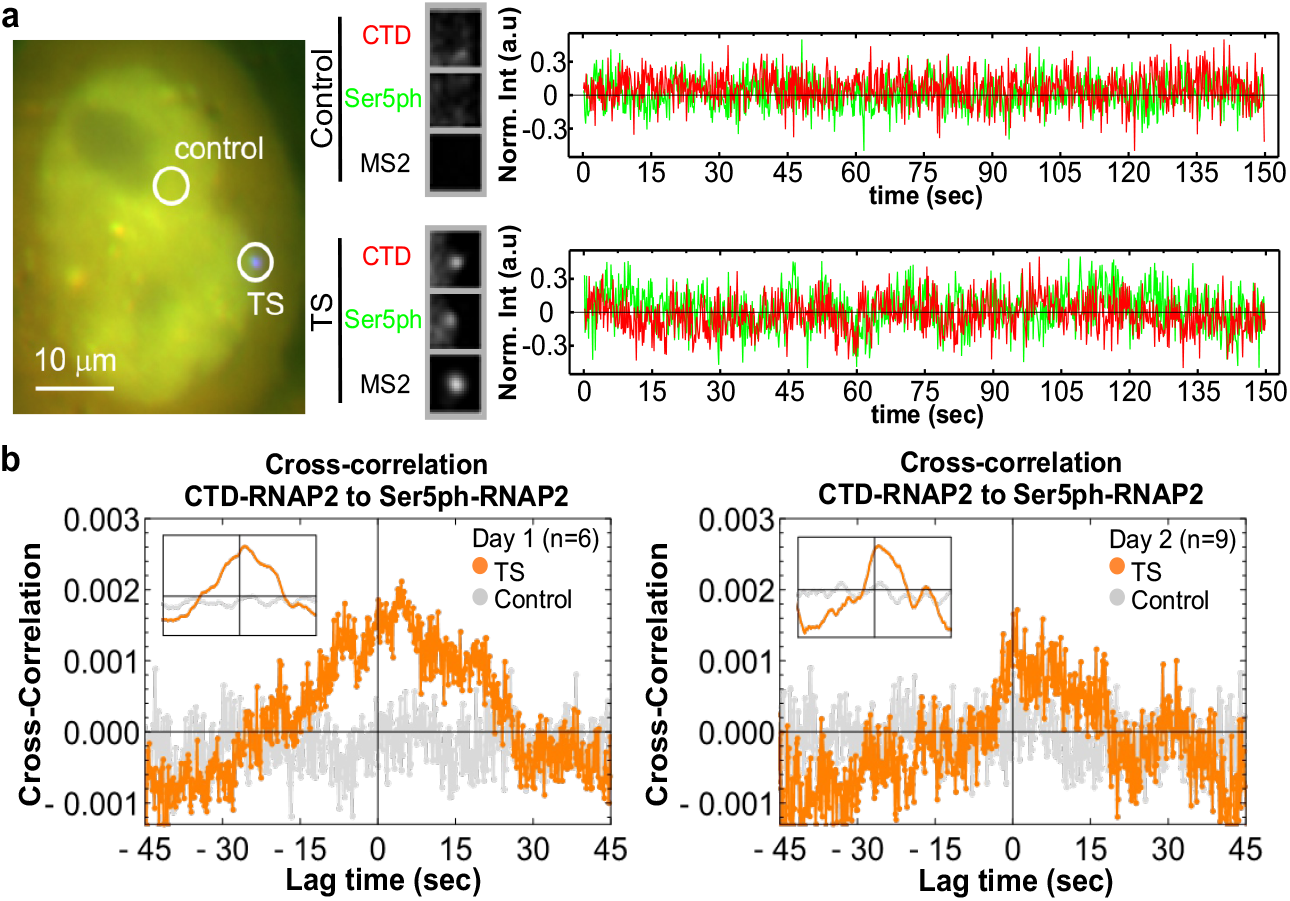
Fast-imaging experiments revealed a 3-6 sec time delay between CTD-RNAP2 and Ser5ph-RNAP2. **(a)** (Left) Exemplary cell for fast imaging (150 ms/frame) for a total of 1000 time points (150 sec) in a single plane. Two positions are highlighted: the HIV-1 TS and a control nonspecific spot. (Center) Crops showing the CTD-(red), Ser5ph-RNAP2 (green), and MS2 mRNA (black) signals at the TS and control positions within the exemplary cell. (Right) Normalized intensity at the TS (bottom) and control (top) positions over time from the exemplary cell for CTD-RNAP2 (red), Ser5ph-RNAP2 (green). **(b)** Measured cross-correlation function *CC*(τ) between CTD-RNAP2 and Ser5ph-RNAP2 at the TS (orange circles) and control (gray circles) positions separated by experimental day. The inset shows a 50-frame rolling average to more easily identify the peak time delay between the two signals. In both cases, the cross-correlation peaks at a lag time of roughly 3-6 seconds.

**Sup. Fig. 6:**
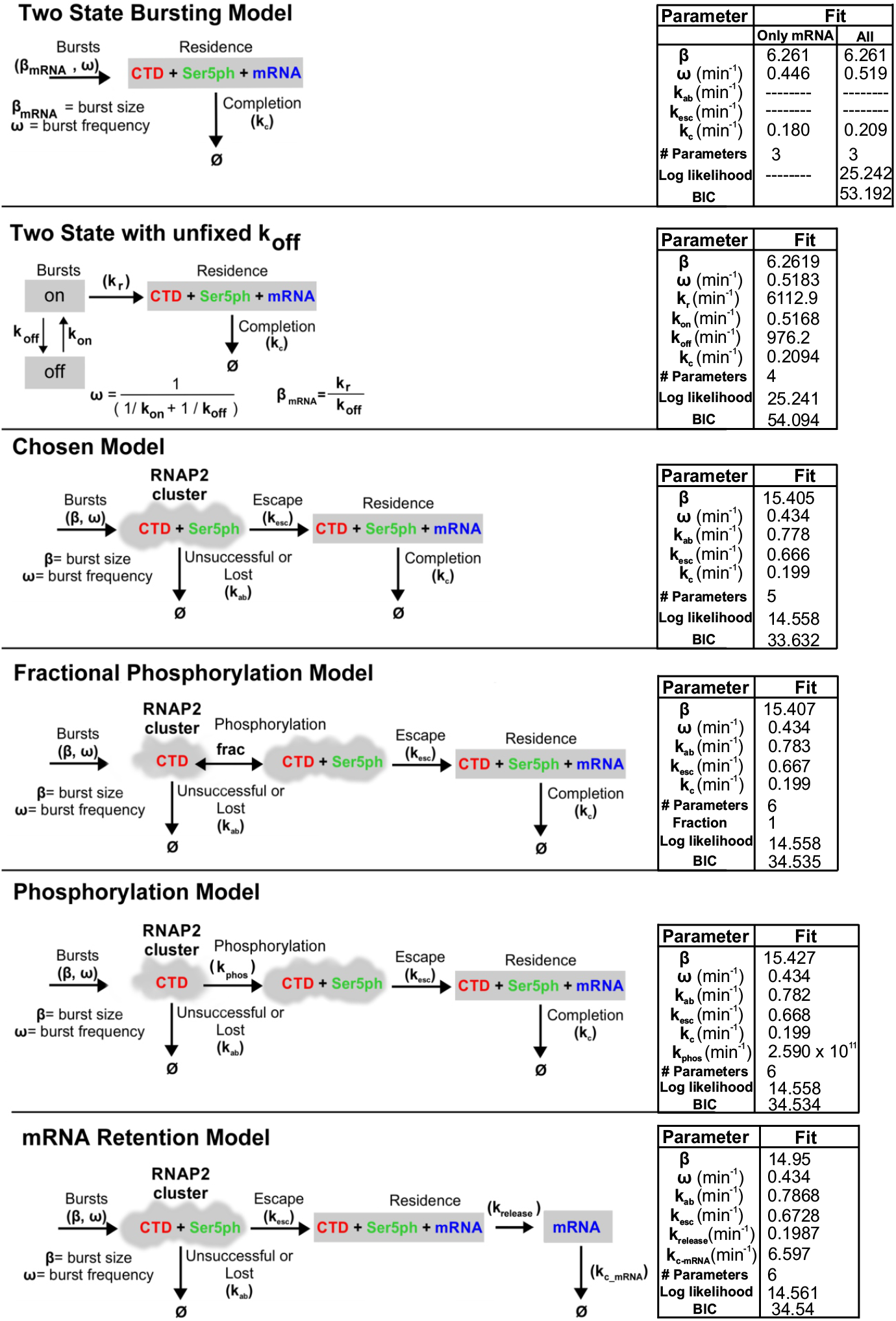
Mathematical models tested. In the simplest “Two-state Bursting Model”, RNAP2 comes in with all 3 signals in a bursting fashion and leaves with a rate *k_c_.* “The Two-State Bursting Model *k_off_* unfixed” is the same as the previous model, however with the parameter *k_off_* not fixed at 1000 and allowed to optimize. The “Chosen Model” is the model described in the main text and in Fig. 3. This model was selected as it fits the experimental data well with a minimum amount of parameters (lowest Bayes Information Criterion, BIC, of tested models). The “Fractional Phosphorylation Model” contains an extra parameter compared to the Chosen Model, frac. Here, frac represents the fraction of unescaped RNAP2 with Ser5ph. Frac is analogous to the ratio of two rates with timescales much faster than the rest of the model: CTD gaining Ser5ph and CTD+Ser5ph losing Ser5ph. In the “Phosphorylation Model”, RNAP2 binds the promoter and then becomes Ser5ph phosphorylated at a rate of *k_phos_*. With this model, RNAP2 requires the Ser5ph signal to escape and transcribe to completion. The “mRNA Retention Model” allows mRNA to remain at the transcription site after RNAP2 completes transcription. All RNAP2 with Ser5ph leave with rate *k_release_* and mRNA are retained at the TS. The mRNA then leave at a rate of *k_c-mRNA_*. Next to each model, the estimated parameters, maximum likelihood, and BIC are shown.

**Sup. Fig. 7:**
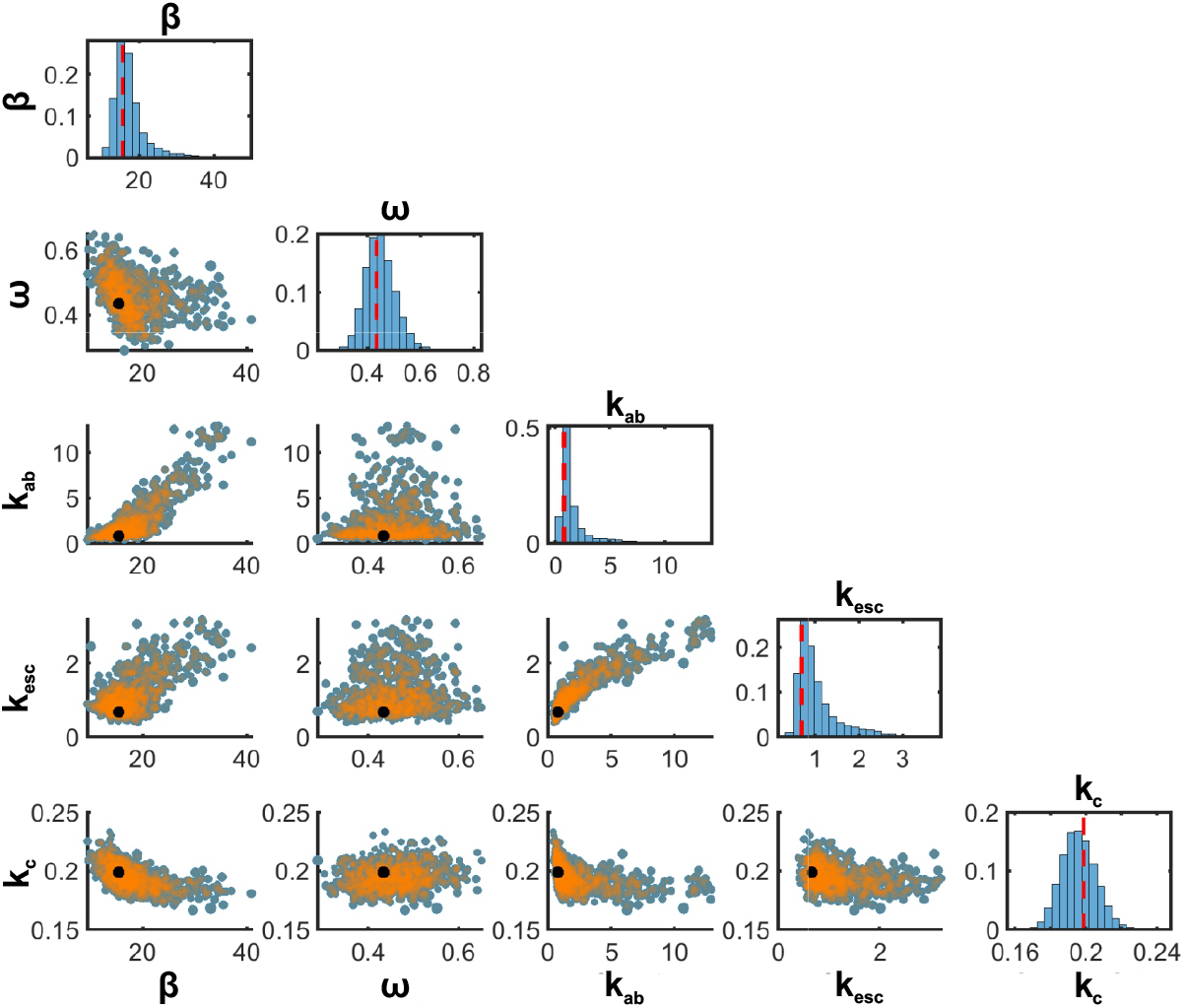
Parameter sensitivity analysis. Metropolis-Hastings algorithm was run to determine posterior uncertainty of model parameters given the experimental data. Plots on the diagonal show the marginal posterior parameter distributions for each parameter (MLE parameter estimate denoted by red dashed line) and off-diagonal plots show the joint posterior parameter distributions for all pairs of parameters (MLE parameter combination denoted by black marker, and high posterior parameters densities are shown in orange).

**Sup. Fig. 8:**
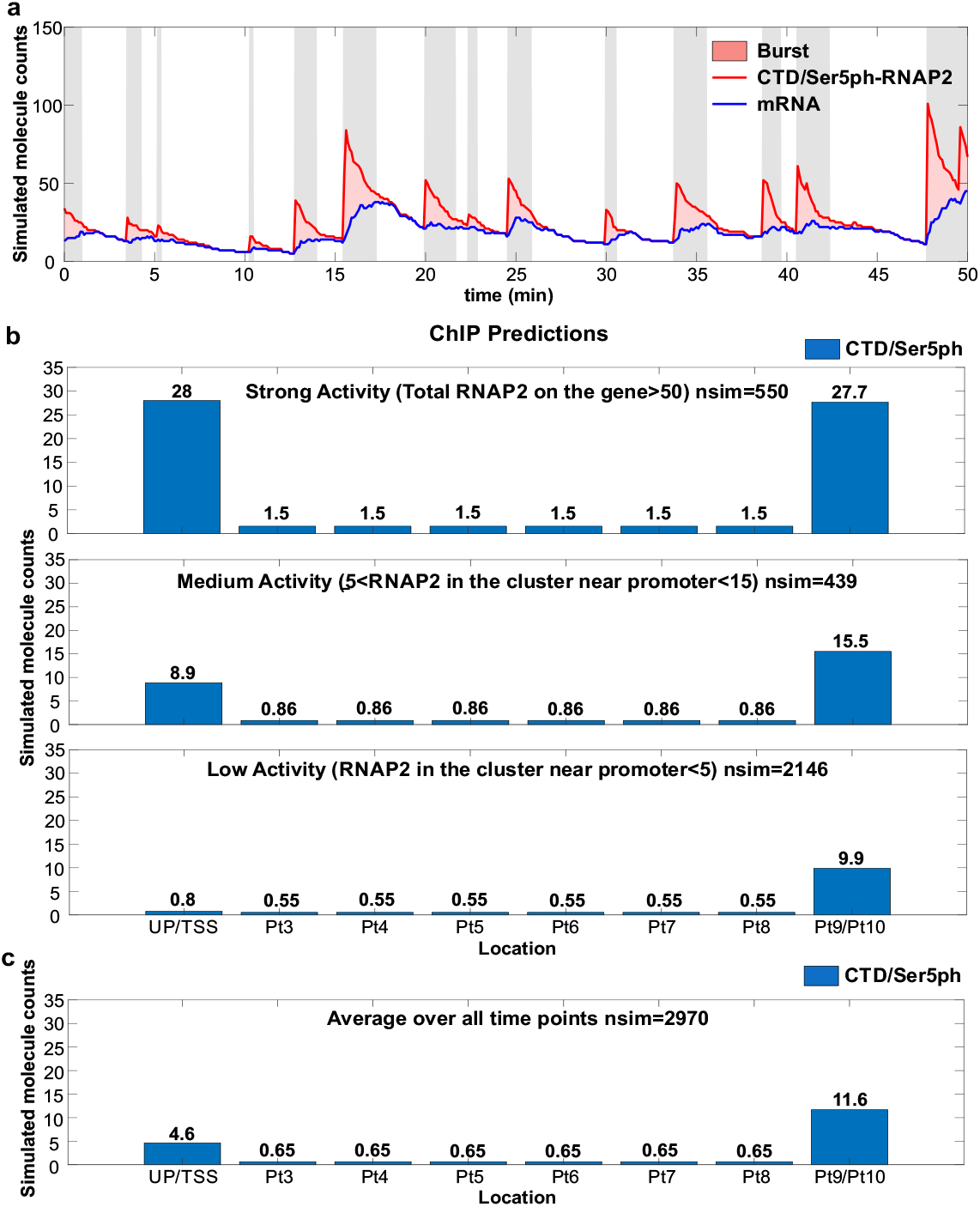
Simulated trajectories and ChIP predictions. **(a)** Stochastic simulation for the number of nascent mRNA per transcription site (blue line), total number of RNAP2 at transcription site (red line), and number of RNAP2 in cluster near transcription site promoter (red shading). Periods with ≥ 10 RNAP2 at the transcription site cluster (gray shading) are classified as ‘ON’ (14.0% of total time); periods with no RNAP2 at the cluster are classified as ‘OFF’ (42.9% of time); and periods with intermediate levels of RNAP2 in the cluster are classified as ‘transient’ (43.1% of time). Note that for clarity these simulations do no include the experimental shot noise used to simulate actual measurements (as in Fig. 3f, for example). **(b)** Simulated ChIP data as predicted using the model for: (Top; Strong Activity) average spot during an ON period; (Middle; Medium Activity) average spot during a transient period; and (Bottom; Low Activity) average spot during an OFF period. Each stochastic simulation was run for 120,000 minutes and sampled at 40 minute intervals to ensure de-correlated points. To estimate RNAP2 loading at the inner bins, an elongation rate of 4.1 kb/min was assumed and used to get the fraction of time spent elongating versus processing of the total RNAP2 residence time. This fraction of elongation time was then distributed from the final bin uniformly to the middle bins and is represented by the middle numbers of bins Pt3-8. **(c)** Average simulated RNAP2 ChIP over all times including all ON, OFF and transient periods.

**Sup. Fig. 9:**
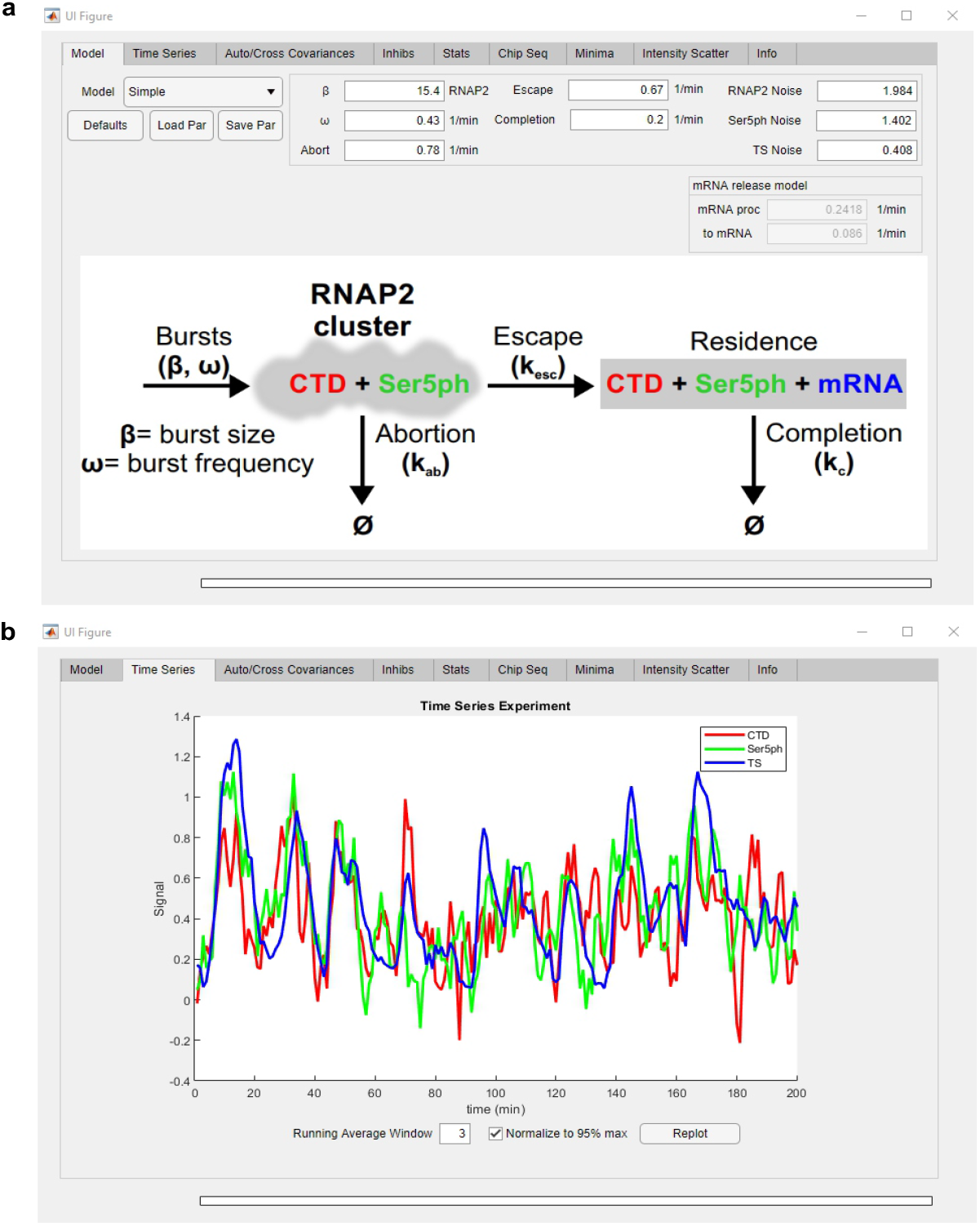
Graphical User Interface (GUI) for the transcription model. To facilitate the simulation of transcription dynamics at a single-copy gene, the model described in the main text has been incorporated into a MATLAB toolbox. **(a)** This graphical user interface (GUI) is divided into eight upper tabs, and input boxes for specification kinetic parameters. The GUI allows the simulation of intensity trajectories in each channel. **(b)** Sample display of simulated intensities normalized to the 95^th^ percentile and running averaged with a window of three time points. The GUI also allows for display of auto-, cross-correlations, predicted minima from the experimental data previously loaded, prediction of ChIP distributions, and perturbed intensity trajectories by blocking different steps of transcription in the model.

**Sup. Fig. 10:**
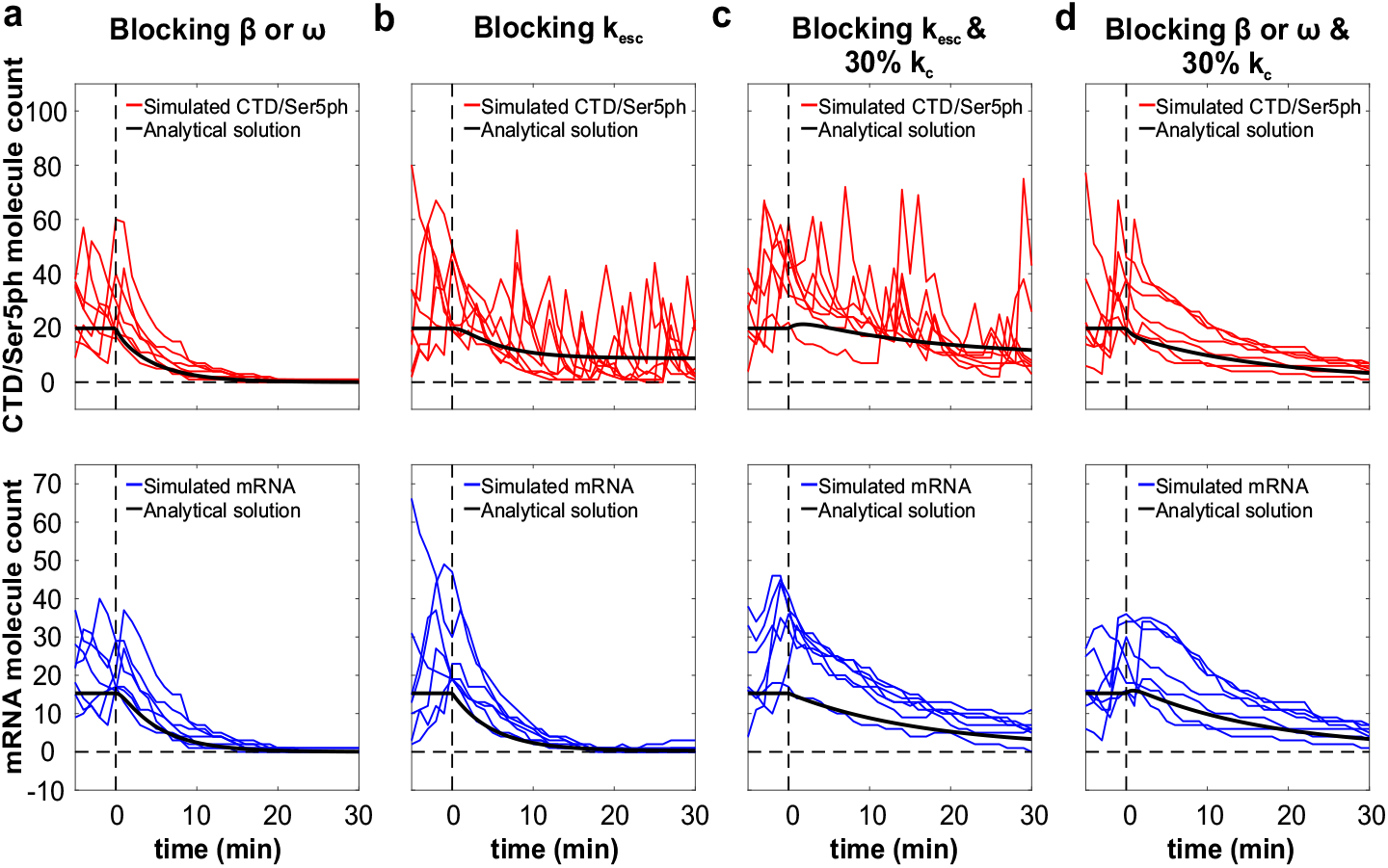
Predicted *CTD/Ser5ph-RNAP2*, and mRNA signals *after* perturbing different steps in the mathematical model. Simulated molecule counts for CTD/Ser5ph-RNAP2 (red, upper panels), and mRNA (blue, bottom panels) after blocking: **(a)** *β* or *ω*, **(b)** k_esc_, **(c)** k_esc_ and 30% k_c_, and **(d)** *β* or *ω* and 30% k_c_, and their respective analytical solution in each plot (black curve). Simulated trajectories with mRNA molecule counts above the analytical solution at time of inhibition are shown with colored lines. This was done to simulate the experimental procedure of choosing transcription sites at the beginning of an experiment where all three signals could be seen. Blocking is defined as multiplying the best fit parameter by 0.01 (99% reduction), similarly blocking 30% refers to multiplying the best fit parameter by 0.3 (70% reduction). For blocking *β* and *ω*, *k_off_* was defined by setting *k_off_* to 1000, effectively turning off bursting dynamics.)

**Table S1:**
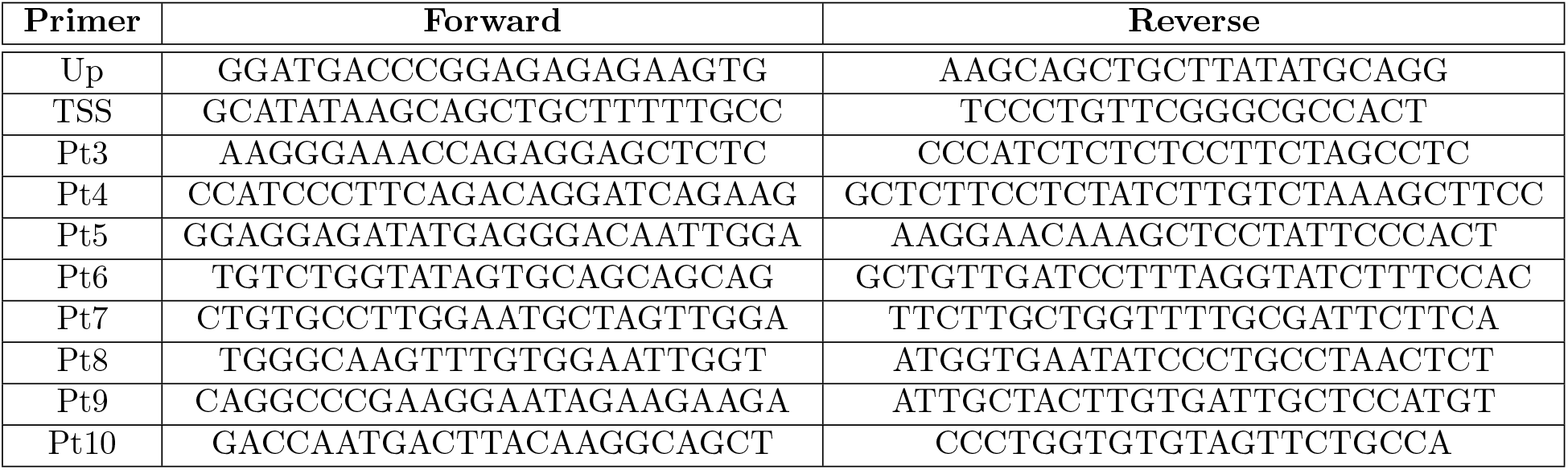
List of primers for ChlP-qPCR as shown in Figure 1. All primer sequences are 5’ to 3’.

***Supplementary Movie 1.*** Dynamics of the cell presented in Fig. 2a, exhibiting multiple cycles of transcription. Maximum projection of a 13 z-stack three-color movie showing an exemplary H-128 cell, in which mRNA (blue), and the RNAP2 Fabs targeting the CTD-RNAP2 (red) and Ser5ph-RNAP2 (green) at the transcription site (within dashed-white circle) of the HIV-1 reporter gene are co-localized. Raw and bandpass filtered crops for each signal and merge (white) over time are shown in the insets. Images were acquired every 1 min for a total of 200 min, shown here as a moving average (over three time points). At select time points, various signals can be seen in the absence of the others, confirming no fluorescence bleed through and demonstrating signals are not perfectly synchronous. For example, t=30-40 min shows a strong burst of all three signals, while t=162-165 min shows CTD-RNAP2 without Ser5ph-RNAP2, t=51-54 min shows Ser5ph-RNAP2 without CTD-RNAP2, and t=149-154 min shows mRNA without the other RNAP2 signals. Scale bar, 10 *μ*m.

***Supplementary Movie 2.*** Dynamics of the cell presented in Fig. 4a, before and after the addition of 5 *μ*M of TPL. Maximum projection of a 13 z-stack three-color movie showing an exemplary H-128 cell co-localizing mRNA (blue), and the RNAP2 Fabs targeting the CTD-RNAP2 (red) and Ser5ph-RNAP2 (green) at the transcription site of the HIV-1 reporter gene before TPL application. Images were acquired every 1 min for a total of 35 min, shown here as a moving average (over three time points). All signals quickly disappeared within 10 min of exposure to TPL. Scale bar, 10 *μ*m.

## References

1. Buratowski, S. Progression through the RNA polymerase II CTD cycle. Mol. Cell 36, 541–546 (2009).

2. Schüller, R. et al. Heptad-specific phosphorylation of RNA polymerase II CTD. Mol. Cell 61, 305–314 (2016).

3. Harlen, K. M. & Churchman, L. S. The code and beyond: transcription regulation by the RNA polymerase II carboxy-terminal domain. Nat Rev Mol Cell Biol. 18, 263–273 (2017).

4. Cramer, P. Organization and regulation of gene transcription. Nature 573, 45–54 (2019).

5. Cissé, I. et al. Real-Time Dynamics of RNA Polymerase II Clustering in Live Human Cells. Science 341, 664–667 (2013).

6. Cho, WK. et al. RNA Polymerase II cluster dynamics predict mRNA output in living cells. Elife 5, e13617-1-31 (2016).

7. Boehning, M. et al. RNA polymerase II clustering through carboxy-terminal domain phase separation. Nat. Struct. Mol. Biol. 25, 833–840 (2018).

8. Lu, H. et al. Phase-separation mechanism for C-terminal hyperphosphorylation of RNA polymerase II. Nature 558, 318–323 (2018).

9. Nagashima, R. et al. Single nucleosome imaging reveals loose genome chromatin networks via active RNA polymerase II. J. Cell Biol. 218, 1511–1530 (2019).

10. Sabari, B. R. et al. Coactivator condensation at super-enhancers links phase separation and gene control. Science 361, eaar3958-1-11 (2018).

11. Heidemann, M., Hintermair, C., Voß, K. & Eick, D. Dynamic phosphorylation patterns of RNA polymerase II CTD during transcription. BBA Gene Regulatory Mechanisms 1829, 5562 (2013).

12. Coulon, A., Chow, C. C., Singer, R. H. & Larson, D. R. Eukaryotic transcriptional dynamics: from single molecules to cell populations. Nat. Rev. Genet. 14, 572–84 (2013).

13. Tokunaga, M., Imamoto, N., & Sakata-Sogawa, K. Highly inclined thin illumination enables clear single-molecule imaging in cells. Nat. Methods 5, 1–7 (2008).

14. Mazza, D., Abernathy, A., Golob, N., Morisaki, T. & McNally, J. G. A benchmark for chromatin binding measurements in live cells. Nucleic acids res. 40, e119-1-13 (2012).

15. Chen, BC. et al. Lattice light-sheet microscopy: imaging molecules to embryos at high spatiotemporal resolution. Science 346, 12579981–12 (2014).

16. Chen, J. et al. Single-molecule dynamics of enhanceosome assembly in embryonic stem cells. Cell 156, 1274–1285 (2014).

17. Li, J. et al. Single-Molecule Nanoscopy Elucidates RNA Polymerase II Transcription at Single Genes in Live Cells. Cell 178, 1–16 (2019).

18. Steurer, B. et al. Live-cell analysis of endogenous GFP-RPB1 uncovers rapid turnover of initiating and promoter-paused RNA Polymerase II. PNAS 115, E4368–E4376 (2018).

19. Bertrand, E. et al. Localization of ASH1 mRNA Particles in Living Yeast. Mol. Cell 2, 437445 (1998).

20. Tantale, K. et al. A single-molecule view of transcription reveals convoys of RNA polymerases and multi-scale bursting. Nat. Commun. 7, 1–14 (2016).

21. Larson, D., Zenklusen, D., Wu, B., Chao, J. & Singer, R. H. Real-time observation of transcription initiation and elongation on an endogenous yeast gene. Science 332, 475–8 (2011).

22. Hayashi-Takanaka, Y., Yamagata, K., Nozaki, N. & Kimura, H. Visualizing histone modifications in living cells: Spatiotemporal dynamics of H3 phosphorylation during interphase. J. Cell Biol. 187, 781–790 (2009).

23. Kimura, H., Hayashi-Takanaka, Y., Stasevich, T. J., & Sato, Y. Visualizing posttranslational and epigenetic modifications of endogenous proteins in vivo. Histochem. Cell Biol. 144, 101109 (2015).

24. Lyon, K. & Stasevich T. J. Imaging translational and post-translational gene regulatory dynamics in living cells with antibody-based probes. Trends in Genetics 33, 322–335 (2017).

25. Conic, S. et al. Imaging of native transcription factors and histone phosphorylation at high resolution in live cells. J. Cell Biol. 217, 1537–1552 (2018).

26. Sato, Y. et al. Histone H3K27 acetylation precedes active transcription during zebrafish zygotic genome activation as revealed by live-cell analysis. Development 146, 1–10 (2019).

27. Stasevich, T. J. et al. Regulation of RNA polymerase II activation by histone acetylation in single living cells. Nature 516, 272–275 (2014).

28. Taube, R., Lin, X., Irwin, D., Fujinaga, K. & Peterlin, B. M. P-TEFb regulation of transcription termination factor Xrn2 revealed by a chemical genetic screen for Cdk9 substrates. Mol. Cell. Biol. 22, 321–331 (2002).

29. Hayashi-Takanaka, Y. et al. Tracking epigenetic histone modifications in single cells using Fab-based live endogenous modification labeling. Nucleic Acids Res. 39, 6475–6488 (2011).

30. Nojima, T. et al. Mammalian NET-Seq Reveals Genome-wide Nascent Transcription Coupled to RNA Processing. Cell 161, 526–540 (2015).

31. Cho, WK. et al. Super-resolution imaging of fluorescently labeled, endogenous RNA Polymerase II in living cells with CRISPR/Cas9-mediated gene editing. Sci. Rep. 6, 1–8 (2016).

32. Zaborowska, J., Egloff, S., & Murphy, S. The pol II CTD: new twists in the tail. Nat. Struct. Mol. Biol. 23, 771–777 (2016).

33. Morisaki, T. et al. Real-time quantification of single RNA translation dynamics in living cells. Science 352, 1425–9 (2016).

34. Coulon, A. et al. Kinetic competition during the transcription cycle results in stochastic RNA processing. Elife 3, 1–22 (2014).

35. Coulon, A. & Larson, D.R. Meth. Enzymol. Fluctuations Analysis: Dissecting Transcriptional Kinetics with Signal Theory, chap. 7 (New York, Academic Press, New York, 2016).

36. Bacia, K., Kim, S. & Schwille S. Fluorescence cross-correlation spectroscopy in living cells. Nat. Methods 3, 83–89 (2006).

37. Müller, F. et al. FISH-quant: automatic counting of transcripts in 3D FISH images. Nat. Methods 10, 277–278 (2013).

38. Munsky, B., Neuert, G. & Van Oudenaarden, A. Using Gene Expression Noise to Understand Gene Regulation. Science 336, 183–187 (2012).

39. Kumar, N., Singh, A., Kulkarni, R. V. Transcriptional Bursting in Gene Expression:Analytical Results for General Stochastic Models. PLoS Comput. Biol 11, e1004292 (2015).

40. Brody, Y. et al. The In Vivo Kinetics of RNA Polymerase II Elongation during Co-Transcriptional Splicing. PLOS Biol. 9, e1000573 (2011).

41. Titov, D. V. et al. XPB, a subunit of TFIIH, is a target of the natural product triptolide. Nat. Chem. Biol. 7, 182–8 (2011).

42. Wang, Y., Lu, J-J., He, L., & Yu, Q. Triptolide (TPL) Inhibits Global Transcription by Inducing Proteasome-Dependent Degradation of RNA Polymerase II (Pol II). PLoS One 6, e23993 (2011).

43. Kwiatkowski, N. et al. Targeting transcription regulation in cancer with a covalent CDK7 inhibitor. Nature 511, 616–620 (2014).

44. Chao, SH. et al. Flavopiridol Inhibits P-TEFb and Blocks HIV-1 Replication. J. Biol. Chem. 275, 28345–28348 (2000).

45. Cho, WK. et al. Mediator and RNA polymerase II clusters associate in transcription-dependent condensates. Science 361, 412–415 (2018).

46. Edelman, L. B. & Fraser, P. Transcription factories: genetic programming in three dimensions. Curr. Opin. Genet. Dev. 22, 110–114 (2012).

47. Feuerborn, A. & Cook, P. R. Why the activity of a gene depends on its neighbors. Trends Genet. 31, 483–490 (2015).

48. Tahirov, T. H. et al. Crystal structure of HIV-1 Tat complexed with human P-TEFb. Nature 465, 747–751 (2010).

49. Darzacq, X. et al. In vivo dynamics of RNA polymerase II transcription. Nat. Struct. Mol. Biol. 14, 796–806 (2007).

50. Guo, Y. E., et al. Pol II phosphorylation regulates a switch between transcriptional and splicing condensates. Nature 572, 543–548 (2019).

51. Barboric, M. & Peterlin, B. M. A new paradigm in eukaryotic biology: HIV Tat and the control of transcriptional elongation. PLoS biology 3, 200–203 (2005).

52. Lionnet, T. & Singer, R. H. Transcription goes digital. EMBO Rep. 13, 313–321 (2012).

53. Core, L., & Adelman, K. Promoter-proximal pausing of RNA polymerase II: a nexus of gene regulation. Genes Dev. 33, 960–982 (2019).

54. Adelman, K. & Lis, J. Promoter-proximal pausing of RNA polymerase II: emerging roles in metazoans. Nat Rev Genet 13, 720–731 (2012).

55. Gu, B., et al. Transcription-coupled changes in nuclear mobility of mammalian cis-regulatory elements. Science 359, 1050–1055 (2018).

56. Straight, A., Belmont, A. S., Robinett, C. C., & Murray, A. W. GFP tagging of budding yeast chromosomes reveals that protein–protein interactions can mediate sister chromatid cohesion. Curr. Biol. 6, 1599–1608 (1996).

57. Viollier, P. et al. Rapid and sequential movement of individual chromosomal loci to specific subcellular locations during bacterial DNA replication. PNAS 101, 9257–9262 (2004).

58. Ochiai, H., Sugawara, T. & Yamamoto, T. Simultaneous live imaging of the transcription and nuclear position of specific genes. Nucleic Acids Res. 43, e127-1-12 (2015).

59. Mariame, B. et al. Real-time visualization and quantification of human Cytomegalovirus replication in living cells using the ANCHOR DNA labeling technology. J. Virol. 92, e00571–18 (2018).

60. Takei, Y. et al. Multiplexed dynamic imaging of genomic loci by combined CRISPR imaging and DNA sequential FISH. Biophys. J. 112, 1773–1776 (2017).

61. Deng, W. et al. CASFISH: CRISPR/Cas9-mediated in situ labeling of genomic loci in fixed cells. PNAS 112, 1870–11875 (2015).

62. Sato, Y. et al. Genetically encoded system to track histone modification in vivo. Sci. Rep. 3, 1–7 (2013).

63. Kimura, H., Tao, Y., Roeder, R.G., Cook, P.R. Quantitation of RNA Polymerase II and Its Transcription Factors in an HeLa Cell: Little Soluble Holoenzyme but Significant Amounts of Polymerases Attached to the Nuclear Substructure. Mol. Cell. Biol. 19, 5383–5392 (1999).

64. Rothbauer, u. et al. Targeting and tracing antigens in live cells with fluorescent nanobodies. Nat Methods 3, 887–889 (2006).

65. Kimura, H., Hayashi-Takanaka, Y., Goto, Y., Takizawa, N., & Nozald, N. The organization of histone H3 modifications as revealed by a panel of specific monoclonal antibodies. Cell Struc. Funct. 33, 61–73 (2008).

66. McNeil, P. L. & Warder, E. Glass beads load macromolecules into living cells. J. Cell Sci. 88, 669–78 (1987).

67. Manders, E. M. M., Kimura, H. & Cook, P. R. Direct imaging of DNA in living cells reveals the dynamics of chromosome formation. J. Cell Biol. 144, 813–821 (1999).

68. Edelstein, A. et al. Advanced methods of microscope control using microManager software. J. Biol. Methods. 1, 1–18 (2014).

69. Carlini, L. et al. Reduced dyes enhance single-molecule localization density for live superresolution imaging. ChemPhysChem 15, 750–755 (2014).

70. Schindelin, J. et al. Fiji: an open-source platform for biological-image analysis. Nat. Methods 9, 676–82 (2012).

71. Aguilera, LU. et al. Computational design and interpretation of single-RNA translation experiments. PLoS Comput Biol. 15, 1–27 (2019).

